# The proton motive force determines *Escherichia coli*’s robustness to extracellular pH

**DOI:** 10.1101/2021.11.19.469321

**Authors:** Guillaume Terradot, Ekaterina Krasnopeeva, Peter S. Swain, Teuta Pilizota

## Abstract

Maintaining intracellular homeostases is a hallmark of life, and key physiological variables, such as cytoplasmic pH, osmotic pressure, and proton motive force (PMF), are typically interdependent. Using a mathematical model, we argue that near neutral pH homeostasis implies that cells must export ions other than protons to generate physiological electrical potential across their plasma membrane. For *Escherichia coli*, proton:ion antiporters are the only known cation efflux pumps, and we therefore predict that principal function of antiporters is to generate an out-of-equilibrium plasma membrane potential and so maintain the PMF at the constant levels observed. Consequently, the strength of the PMF determines the range of extracellular pH over which the cell is able to preserve its near neutral cytoplasmic pH, and the non-zero PMF is needed to maintain membrane potential. In support, we concurrently measure the PMF and cytoplasmic pH in single cells and demonstrate both that decreasing the PMF’s strength impairs *E. coli*’s ability to maintain its pH and that artificially collapsing the PMF destroys the out-of-equilibrium plasma membrane potential. We further predict the observed ranges of extracellular pH for which three of *E. coli*’s antiporters are expressed, through defining their cost by the rate at which they divert protons from being imported to generate ATP. Taken together, our results suggest a new perspective on bacterial electrophysiology, where cells regulate the plasma membrane potential to maintain

---

Internally, living cells differ from their environment. They typically maintain intracellular properties within a restricted range in the phenomenon of homeostasis. For example, cytoplasmic, or internal, pH (pH_*i*_) is kept close to neutral in organisms across all kingdoms of life [1, 2], likely because pH_*i*_ affects the stability and function of proteins [3].

Neutralophilic bacteria, such as *Escherichia coli*, maintain an intracellular pH close to 7.5 over a range of extracellular pH (pH_*e*_) [4, 5, 6]. For example, after a sudden shift in pH_*e*_ from near neutral to acidic values, *E. coli*’s pH_*i*_ initially follows the change in pH_*e*_, but in some experiments recovers to ∼7.5 [7, 8, 9] and in some more recent ones does not (Fig. 1A) [10, 11, 12]. Explanations offered for this discrepancy include strain- to-strain differences and inaccuracies in calibrating and usage of pH sensors [10].

**Fig. 1.**
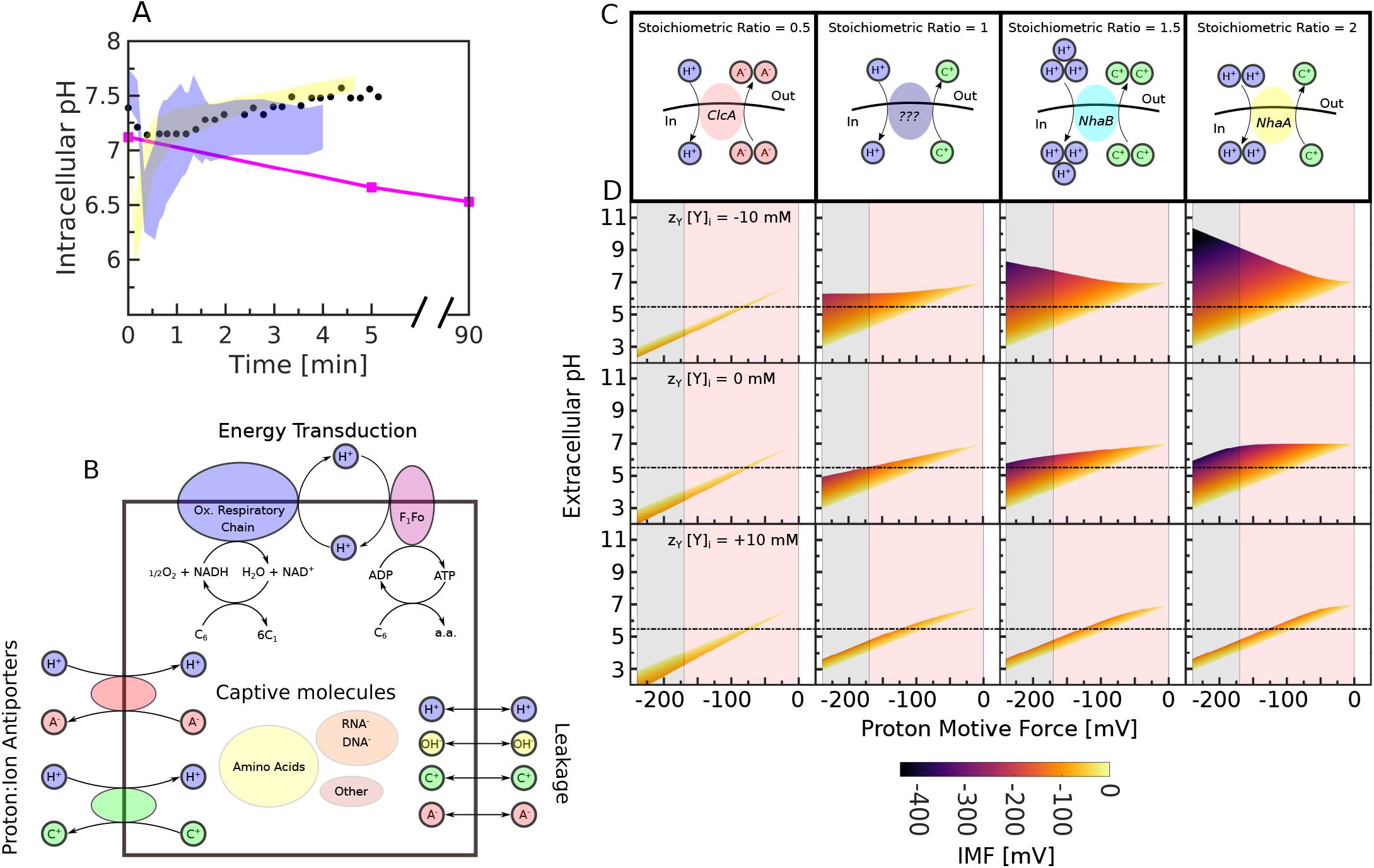
The mathematical model of *E. coli’s* electrophysiology predicts that the pH_*e*_ range over which cells can maintain a neutral pH_*i*_ is directly related to the PMF. (A) We reproduce contradictory experimental results on the response of pH_*i*_ in *E. coli* to changes in pH_*e*_. Pink squares are for a pH_*e*_ shift from 7.2 to 5.6 [10]. Black dots are population averages for a pH_*e*_ shifted from 7.5 to 5.5 [7], and the yellow [8] and blue [9] shaded areas show the range of pH_*i*_ measured in single cells for the same shift in pH_*e*_. (B) A simplified model of *E. coli*’s electrophysiology. We distinguish two classes of chemical species contributing to the membrane potential — small ions, which both leak and are pumped across the membrane, and captive molecules, which cannot cross the membrane. Only two cations, protons and a representative cation C^+^, and two anions, hydroxide ions and an anion A^−^, are included. Cells generate the PMF by either the electron transport chain or the F_1_ F_*o*_ ATPase. We fix the concentrations of NADH, NAD^+^, ATP, and ADP, and of extracellular ions. Proton-ion antiporters exchange protons for either C^+^ or A^−^ and generate ionic motive forces. (C) We consider four antiporters. Their stoichiometric ratio is the ratio of the number of protons to the number of ions exchanged. (D) We plot the possible steady-state solutions for each antiporter for a given PMF between -240 mV and 0 mV, a pH_*i*_ of 7, and pH_*e*_ between 2 and 12. Extracellular [CA]_0_ = 100 mM, and we consider three values of the total captive charge, *z*_*Y*_ *Y*_*i*_. The horizontal black line marks pH_*e*_= 5.5. Grey and red shading shows values of the PMF expected for respiration and fermentation. The colour scale indicates the ionic motive force (IMF).

However, the homeostasis of pH is rarely studied in the context of all the other physiological variables that pH influences and the cell regulates. For example, the electrical potential across the membrane – the membrane voltage Δ*ψ* – is generated by the charge accumulated at the membrane, which includes protons. This Δ*ψ* and the difference between the intracellular and extracellular pH (ΔpH) determine the proton motive force (PMF), the electrochemical gradient of protons. More generally, the electrochemical gradients of all the other ions present in the cell – the ion motive forces (IMFs) – are influenced too by the Δ*ψ* and so indirectly by protons. Protons can also affect the osmotic pressure of the cell (Π), which depends on the difference between the extracellular and intracellular concentrations of all solutes [13].

The intertwined nature of these physiological variables suggests that we should study their maintenance together even if we wish to understand the homeostasis of one. To do so, we built a mathematical model of bacterial electrophysiology. We show mathematically that the low quantities of intracellular protons necessitated by a near neutral pH_*i*_ imply that these protons contribute little directly to the membrane potential. We next demonstrate that cells build a membrane potential using proton:ion antiporters to drive intracellular ion concentrations away from equilibrium. As a consequence, we predict that the PMF determines the cell’s ability to maintain membrane potential, as well as near neutral pH_*i*_ as pH_*e*_ changes. We measured membrane potential at zero PMF and pH_*i*_ during shifts in extracellular pH at two different magnitudes of the PMF. Agreeing with the model, we find that cells cannot maintain membrane potential in absence of the PMF, and maintain pH_*i*_ at a given pH_*e*_ only if the absolute magnitude of the PMF is sufficiently high. Together our results suggest that the primary role of proton:ion antiporters in *E. coli* is to modulate membrane potential to maintain the PMF. This is in contrast with their previously assumed role of direct pH_*i*_ regulation by importing or exporting protons depending on the pH_*i*_. While we do not exclude a (different kind of) direct pH_*i*_ regulation, we conclude that it must be constrained by the antiporters’ role in modulating membrane potential.

## Results

### A mathematical model provides an integrated view of bacterial electrophysiology

To describe as simply as possible the intertwined physiological variables, we developed a minimal mathematical model of bacterial electrophysiology, building on our previous work [14]. We consider pH_*i*_, which we assume is kept near neutral, the electrochemical gradients of ions — the IMFs, and osmotic pressure.

For a single ion (*x*), the electrochemical potential is the free energy required to move an extracellular ion into the cell measured in volts. Mathematically, this potential is defined as

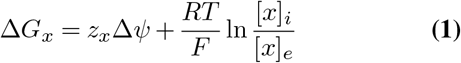

with an intracellular concentration of [*x*_*i*_] and an extracellular concentration of [*x*_*e*_]. *z*_*x*_ is the ion’s valency, *R* is the gas constant, *T* is temperature, *F* is Faraday’s constant, and Δ*ψ* is the membrane potential. We will use Δ*G*_*x*_ interchangeably with the IMF.

A build-up of electrical potential across the membrane is enabled by its hydrophobicity, and, following others [15, 16], we assume that the charge accumulated in close proximity to the membrane is equivalent to the surface charge on a parallel-plate capacitor. Writing the membrane’s capacitance as *C*_*m*_, the cell’s volume as *V*, and its surface area as *S* (*Supplementary Information*), we have

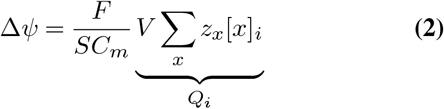

where *Q*_*i*_ is the total intracellular charge. *Q*_*i*_ is only non-zero adjacent to the membrane and is equal and opposite in sign to the total extracellular charge *Q*_*e*_, which is also only non-zero adjacent to the membrane. For simplicity, and without any loss of generality — a rescaling is required otherwise, we assume that the cell has no incompressible volume.

The IMF of particular relevance for *E. coli* is the proton motive force, the PMF or 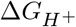. A sufficiently high PMF is not only relevant for transport, but also for ATP synthesis via the F_1_F_*o*_-ATP synthase and for motility [17]:

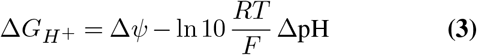

where ΔpH = pH_*i*_ − pH_*e*_.

Lastly, bacteria maintain osmotic pressure within a given range: 0.3 to 3 atm for *E. coli* [18, 19]. Assuming osmotic coefficients equal to one, Π is proportional to the difference in total concentration across the membrane [16, 20]:

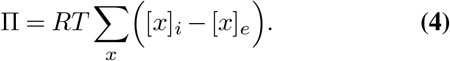

Our model comprises five species: protons, H^+^, hydroxide ions, OH^−^, a monovalent cation, C^+^, a monovalent anion, A^−^, and a lumped species, *Y*, that includes the contribution to Π and Δ*ψ* of all intracellular molecules that are not these four ions (Fig. 1B). These could include charged molecules that are not able to cross the plasma membrane, or that do so with processes we do not model, e.g. free glutamate – the most abundant [21], as well as DNA and RNA. Although we do not specify the anion or the cation, we do assume that the equivalent cation would be uniquely mapped, i.e. either Na^+^ or K^+^.

All ions except captive ions leak across the plasma membrane [16], but H^+^, C^+^ and A^−^ may also be actively transported by antiporters. To describe how the concentration of C^+^ and A^−^ change we write

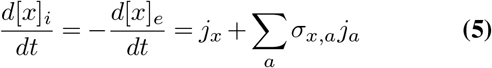

where *j*_*x*_ is the flux from leakage and *j*_*a*_ is the flux from antiporters, each with a stoichiometric coefficient *σ*_*x,a*_. A proton antiporter with stoichiometric ratio of two, for example, imports two protons and exports one cation. For the intracellular concentrations of protons and hydroxide ions, we include the self-ionization of water at a rate *j*_*w*_ (see also *Supplementary Information*):

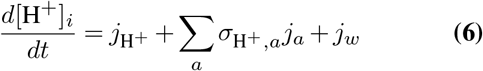

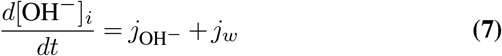

Both the flux from active transport and the leakage depend on the Δ*G* of the reaction [22]:

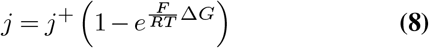

where the transport mechanism, such as the number of intermediate steps involved, determines the rate *j*^+^ [16]. Like others, we assume that *j*^+^ is constant for active transport at steady state [15], and, for leakage, we use an expression that depends on the membrane’s permeability (*Supplementary Information*).

We first note that without active transport, ions will leak across the membrane making Δ*G*_*x*_ = 0. The membrane potential will not vanish, however, because of any captive molecules that are charged. This equilibrium Δ*ψ*, or Donnan potential, cannot reach physiologically levels without generating an excessive Π (Fig. S1). To generate both a physiological Δ*ψ* and Π, cells must therefore actively transport ions.

Second, by considering four possible scenarios of active transport, anions or cations either in or out (Fig. S2), we find that for near neutral pH_*i*_ only exporting cations robustly generates both a physiological Δ*ψ* and Π. The others produce values outside the experimentally observed range (*Supplementary Information*). Exporting anions results in positive Δ*ψ*, making it a viable strategy only in a narrow range of acidic extracellular environments. These conclusions hold if we allow both anions and cations to be pumped simultaneously, Fig. S3.

Third, we model that the active transport of ions is ultimately powered by the cell’s metabolism, which leads to the export of protons by either oxidising NADH or hydrolysing ATP [23]. We refer to these regimes in our model as respirative and fermentative like. However, at a near neutral pH_*i*_ the concentrations of protons and hydroxide ions are too low, at 10^−3^ mM, to contribute substantially to the Δ*ψ*. Even if pH_*i*_ is within the 6-8 range, the extremes for most living cells [1, 10], their contribution is at most *±*3.5 mV (whereas the reported values of Δ*ψ* are around −140 mV [24]). We, therefore, assume that the cell re-imports these protons to export other ions through proton:ion antiporters, because the intra-cellular concentrations of other ions are much less restricted and because no other types of cation (apart from proton) exporters, such as ATP-driven, are known to exist in *E. coli*.

Finally, and following observations [25, 26], we assume that cells maintain a constant, time-invariant value of the PMF (in *Supplementary Information* we determine its physiological limits). To do so at a near neutral pH_*i*_ the cells must control Δ*ψ*, modifying its value in response to changes in pH_*e*_.

### The cell’s PMF determines the range of pH_*e*_ at which it maintains near neutral pH_*i*_

In our model, metabolism powers the export of protons. To generate physiological Δ*ψ*, and thus PFM, cells change the concentrations of other ions only through the action of antiporters.

To understand how the PMF’s value (generated in such a way) influences pH_*i*_, we look at four types of antiportres (Fig. 1C), three of which have identical stochiometric ratios to antiporters found in *E. coli*: ClcA [27], NhaB [28], and NhaA [29]. The fourth, which we depict with question marks, has no known equivalent and exports one cation in exchange for one proton.

Although changes in pH_*e*_ may generate a change in expression of antiporters or changes in their stochiometric ratio, as part of a possible regulatory mechanism [6, 30], we included such effects only implicitly in the value of *j*^+^ in Eq. (8). This value once positive is not critical for much of our analysis.

The reaction powered by the antiporter *a* in Eq. (5) is

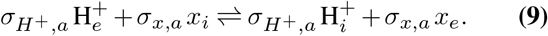

To export *x*, this reaction’s Δ*G* should be negative:

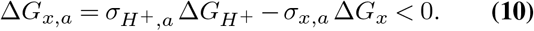

At steady state, the leakage of *x* back across the membrane balances this export, implying that the leakage flux *j*_*x*_ must be positive and, from Eq. (8), have Δ*G*_*x*_ *<* 0. Combining with Eq. (10) implies:

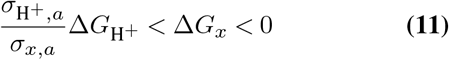

and the IMF, Δ*G*_*x*_, is bounded. The two limits correspond to the antiporter working maximally when Δ*G*_*x*_ is most negative or negligibly when Δ*G*_*x*_ is zero.

For a given antiporter, characterised by its stochiometric ratio, we wish to determine the range of steady states that the antiporter can support. By fixing the concentrations of extracellular ions, pH_*e*_, and the charge and intracellular concentration of captive molecules, and by assuming that the pH_*i*_ is neutral, we specify a steady state by the value of the PMF. If an antiporter can maintain this steady state, then Eq. (11) must hold.

To check if Eq. (11) is valid, we must determine Δ*G*_*x*_. First, we find the membrane potential. Because we have specified pH_*i*_, pH_*e*_ and the PMF, we use the definition of the PMF, Eq. (3). Second, we simplify Eq. (2) to become:

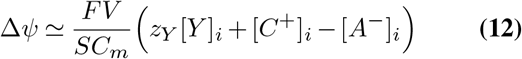

by ignoring the negligible contributions of protons and hydroxide ions. We then use Eq. (1) to include the known extracellular concentrations

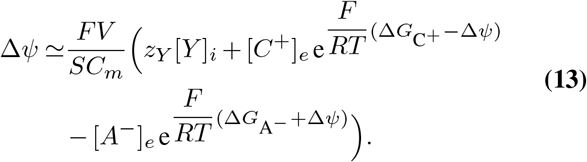

Third, the antiporter pumps only one of these ions. The other leaks across the membrane and must have Δ*G* of zero at steady state. Eq. (13) then reduces to an equation for Δ*G*_*x*_. We solve this equation and evaluate Eq. (11).

The result is that the antiporters with different stochiometries generate steady states with pH_*i*_=7 for different ranges of PMF and pH_*e*_ (Fig. 1D, S4 and S5).

For cation-exporting antiporters, such as NhaA and NhaB, the lower limit of pH_*e*_ that supports a steady state is determined by the upper bound of Eq. (11). When pH_*e*_ decreases below pH_*i*_ for a constant PMF, the cell generates a more positive membrane potential to maintain the PMF from Eq. (3). For cation-exporting antiporters, which work to build negative Δ*ψ* and so negative 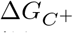, weaker pumping antiporters that allow cations to build up decreasing the magnitude of 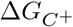 are necessary. At the minimal possible pH_*e*_, the pumping by the antiporters is negligible, and the magnitude of 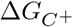 is close to zero. For these antiporters, the upper limit of pH_*e*_ is determined by the lower bound of Eq. (11). When pH_*e*_ increases above pH_*i*_ for a constant PMF, the cell generates a more negative membrane potential to maintain the PMF with stronger pumping cation-exporting antiporter, increasing the magnitude of 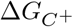. At the maximal possible pH_*e*_, the antiporter builds 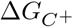 and Δ*ψ* to the maximal magnitude it can, and 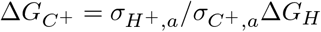. This behaviour will hold too for an electroneutral cation-exporting antiporter.

Alternatively, for ClcA-like antiporters, which use the PMF to generate a more positive Δ*ψ* by exporting anions, it is the lower bound of Eq. (11) that sets the lower limit of pH_*e*_. At this bound, the antiporters increase Δ*ψ* by exporting anions as much as they can, and the absolute 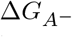 is maximal. The antiporters are importing protons to generate this behaviour even though the extracellular environment is acidic; they do not need to export protons to raise pH_*i*_ directly as previously thought [1, 31]. The upper bound of Eq. (11) sets the upper limit of pH_*e*_. The antiporters generate the more negative Δ*ψ* needed by more weakly exporting anions.

The range of pH_*e*_ over which the cell maintains a steady state with pH_*i*_=7 increases if the PMF is more negative for all the antiporters we tested (Fig. 1D, see also discussion in *Supplementary Information*). A more negative PMF both increases the free energy available to the antiporters, increasing the maximal flux at which they can work and extending the lower bound of Eq. (11), and, from Eq. (3), increases the Δ*ψ* at neutral pH_*e*_, decreasing Δ*G*_*x*_ further away from its maximal value of zero.

We therefore predict that cells can maintain pH_*i*_ over the widest range of pH_*e*_ in the respiratory rather than the fermentative regime because the PMF is then more negative (Fig. 1D). This robustness to pH_*e*_ comes at a cost. A more negative PMF will require the electron transport chain to export protons at a greater rate because the correspondingly more negative membrane potential will increase the import rate of protons by both the antiporters and leakage.

For a hypothetical ATP-driven cation efflux pump — some mammalian cells generate Δ*ψ* using a sodium-potassium ATPase [32], these observations would not hold. As for the antiporters, the Δ*G* of such pumping reaction should be negative and the reaction balanced by leakage at steady state, implying:

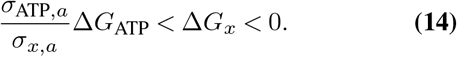

The IMF is now bounded below by the free energy of ATP hydrolysis rather than the PMF (Fig. S6).

### At pH_*e*_ 5.5, *E. coli* maintains pH_*i*_ if the PMF is sufficiently negative

To confirm the PMF’s predicted role on the robustness of pH_*i*_, we should measure PMF and pH_*i*_ simultaneously in individual bacteria while controlling the magnitude of the PMF. We can do so by using the speed of the bacterial flagellar motor to indicate PMF and a ratiometric pH sensor (pHluorin [33]) to measure pH_*i*_ [34, 14, 35].

The flagellar motor is a rotary nano-machine [17], and its speed is proportional to the PMF [36, 37, 24] (Fig. S7). Measuring changes in the motor’s speed is therefore equivalent to measuring changes in the PMF. By attaching a cell to the surface of a cover slip [38] and a plastic bead to a genetically modified filament stub of the flagellum [39, 40, 41, 34, 35], we are able to measure the rotation of the motor labelled with the bead through placing the bead in a heavily attenuated optical trap and performing back-focal-plane interferometry (Fig. 2A) [42, 43]. The strains we use also express cytoplasmic pHluorin [34, 14, 35].

**Fig. 2.**
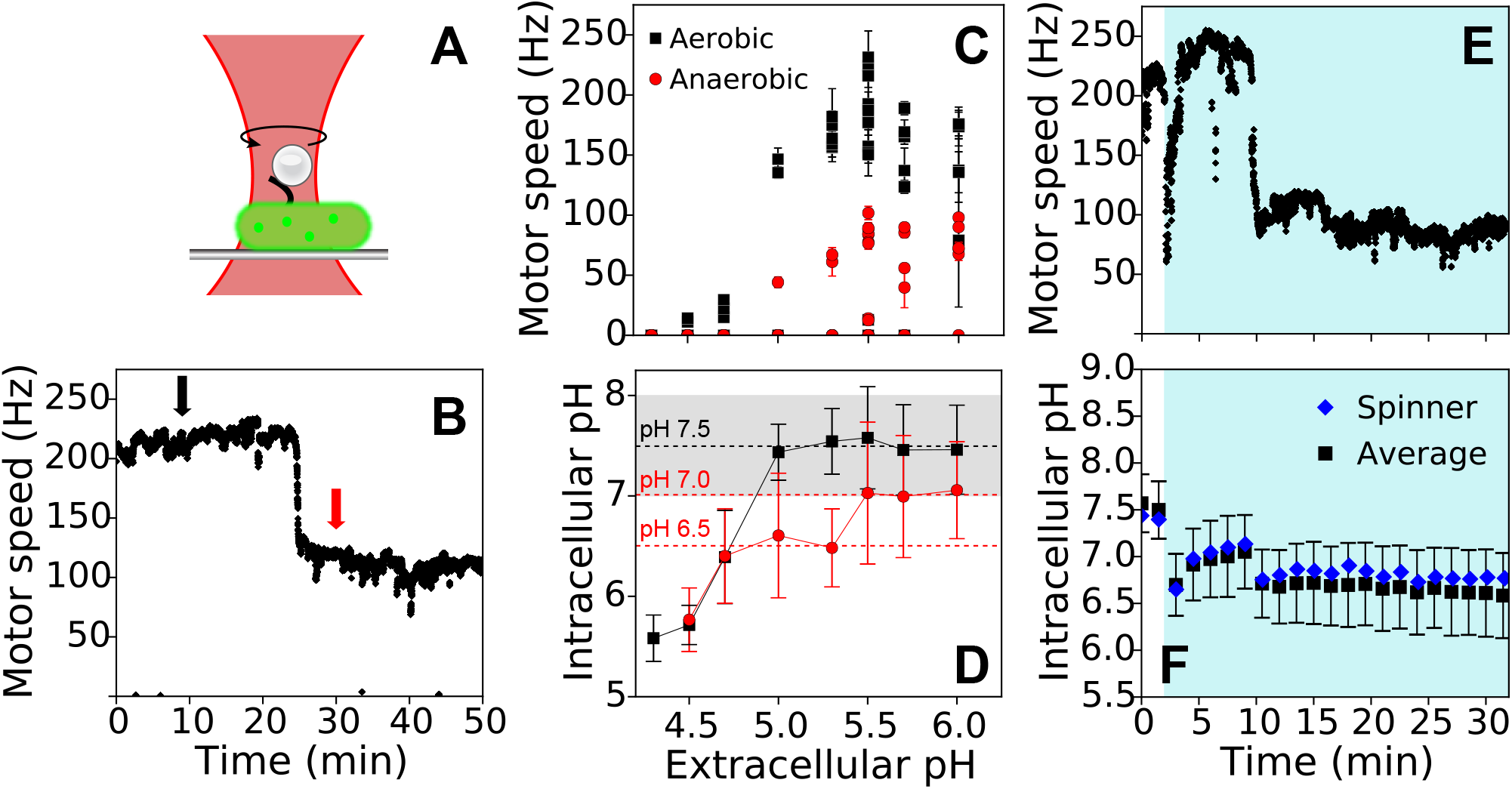
At acidic pH_*e*_, *E. coli* maintains neutral pH_*i*_ if the magnitude of the PMF is sufficiently high. (A) A schematic showing *E. coli* immobilised on a glass coverslip with a polystyrene bead attached to a ‘sticky’ stub of a bacterial flagellar filament. We place a bacterium in the focus of a heavily attenuated laser to detect the position of the bead at high speeds and use the flagellar motor’s rotational speed as a proxy for PMF. The cell’s cytoplasm is in green to illustrate that we concurrently monitor a fluorescent pH sensor. (B) An example of the motor speed from a single cell in a sealed tunnel-slide at pH_*e*_=7.0 in 20 mM glucose. The sharp drop at ∼ 25 min occurs when oxygen is exhausted. The arrows show where we measure ‘aerobic’ (black) and ‘anaerobic’ (red) speeds. (C) The single-cell motor speed depends on pH_*e*_. We calculate speeds as the mean of a *±* 45 s interval around the time points indicated in B. Error bars are standard deviations. (D) pH_*i*_ changes with pH_*e*_. Values of pH_*i*_ are the mean of 90 to 316 cells for the time points indicated in B; error bars are standard deviations. Grey shading shows the region of pH_*i*_ homeostasis, between pH 7 and 8. (E) An example of single-cell motor speed upon shifting to pH_*e*_=5.5. Initially cells are at pH_*e*_=7, and then we shift pH_*e*_ to 5.5 at *t*= 2 min (blue shade) and seal the tunnel-slide. At the point where pH_*e*_ changes, the bead rotated by the motor moves out of the laser’s region of detection, due to flow, giving a loss of signal — the sharp drop in speed. (F) The dynamics of pH_*i*_ depends on the PMF. Blue diamonds show the cell from E; black squares show the mean and the standard deviation for all the cells in the field of view (∼ 30). Blue shading indicates where pH_*e*_ is 5.5.

To change the PMF, we use a sealed tunnel-slide configuration [44, 41, 34]. Cells are kept in a potassium phosphate buffer containing NaCl to match the osmolality of the growth media and glucose. They eventually run out of oxygen, at which point the magnitude of the PMF drops [45] (Fig. 2B). Although the timing of the drop varies between 20 and 30 min and depends on the concentration of cells on the tunnel-slide surface, as found too for a population of swimming cells [45], there is a conserved characteristic step-like shape (Fig. S8).

We change pH_*e*_ from 7.0 to a lower value (4.3 - 6) and observe pH_*i*_, first at a higher absolute PMF that cells maintain in the presence of oxygen and then at lower one when the oxygen in the tunnel-slide runs out (arrows in Fig. 2B). The motor’s speed, and hence the PMF, changes in these aerobic and anaerobic conditions (Fig. 2B & C).

As predicted (Fig. 1D), cells maintain pH_*i*_ over a wider range of pH_*e*_ for the higher magnitude PMF in the aerobic environment compared to the lower magnitude PMF in the anaerobic environment (Fig. 2C & D).

Although the predictions in Fig. 1D are for steady states, experimentally we can also monitor transitions between these steady states. For example, changing pH_*e*_ to 5.5 temporarily acidifies the cytoplasm (Fig. 2F), but pH_*i*_ then recovers towards its initial value if the magnitude of the PMF is kept high by oxygen being present (Fig. 2E). As oxygen falls and the PMF’s magnitude drops, pH_*i*_ falls back to the value seen when pH_*e*_ was first changed. If we alter pH_*e*_ to below 5, we observe no dynamic recovery, and both the magnitude of the PMF and pH_*i*_ drop permanently indicating that the gradient of protons across the membrane has collapsed (Fig. S9).

### A ΔpH requires a PMF if antiporters generate the membrane potential

Another prediction of our model is that cells require a PMF to maintain a ΔpH. When the PMF is zero, ΔpH is proportional to Δ*ψ*, from Eq. (3), but without a PMF the antiporters cannot drive the membrane potential from equilibrium, and so ΔpH should be zero. Fig. 3B shows that for antiporters with stoichiometric ratios of 0.5, 1.5 and 2, Fig. 3A, this is the case. If the cells had an ATP-driven efflux pump, however, then they could generate a membrane potential with no PMF, and ΔpH would not be zero, as shown in Fig. 3B far right. We note that with captive charged molecules, our antiporter model does not predict a Δ*ψ* that is exactly zero because the captive charges mean that the equilibrium Δ*ψ* is not zero.

**Fig. 3.**
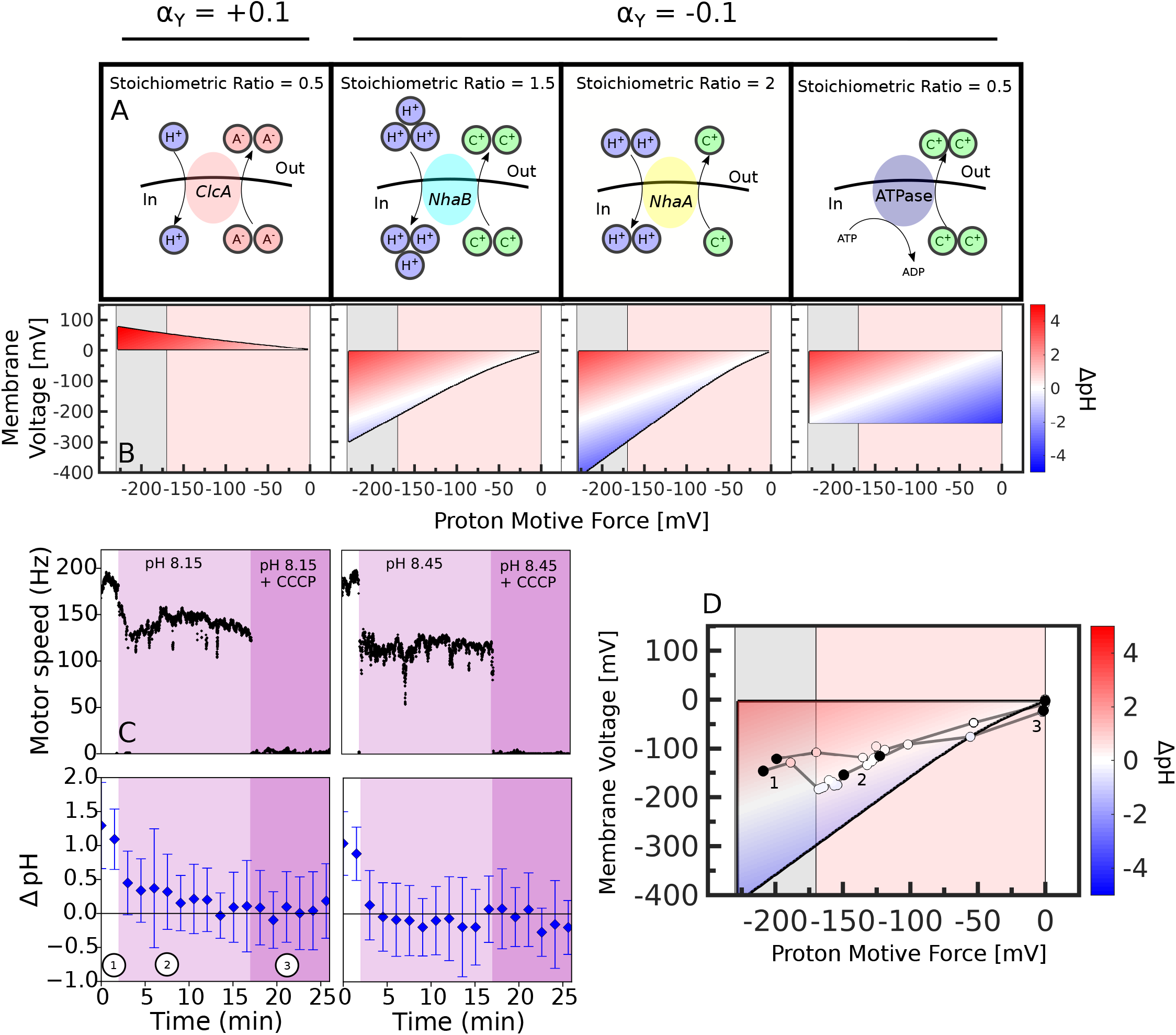
Collapsing the PMF gives results consistent with proton antiporters generating *E. coli*’s membrane potential. (A) Schematics of the ion pumps in the model. We include a hypothetical ATPase with a stoichiometry of one ATP per two cations. (B) Generating Δ*ψ* by antiporters or by an ATPase gives a different ΔpH = pH_*i*_ − pH_*e*_ when the PMF is close to zero. We plot the possible steady-state membrane potentials as a function of the PMF and show ΔpH using the colour scale with pH_*i*_ = 7 (blue is alkaline pH_*e*_; red is acidic). The anionic antiporters maintain a positive potential; the cationic antiporters maintain a negative potential. For computational reasons, we show an approximate Δ*ψ* (with an error of ∼10 mV – Fig. S9). (C) The single-cell motor speed (top panels), reporting on the PMF, and the average ΔpH (bottom panels) upon alkaline shifts to 8.15 (first column) or 8.45 (second column) followed by addition of a protonophore, CCCP, support a Δ*ψ* generated by antiporters. We change pH_*e*_ from 7 (white — annotated 1) to above 8 (pink — annotated 2) and then add CCCP (purple — annotated 3). ΔpH is the mean of ≥30 cells; errors are with standard deviations. (D) The data qualitatively agrees with the model’s prediction of a vanishing membrane potential. Estimating the motor speed from C using a conversion factor of 1.13 mV/Hz (Fig. S7) and the membrane potential from Eq. (3) for the NhaA-like antiporter in B, we plot the estimated membrane potential versus the PMF. The black annotated dots show specific time points in C, and the grey and red shading indicate the respiratory and fermentative regimes.

To collapse the PMF experimentally, we treat cells in the alkaline regime (where the difference between antiporters and a putative ATP-drive efflux pumps is most notable in Fig. 3B) with a commonly used protonophore (Fig. 3C), 100 *μ*M carbonyl cyanide m-chlorophenylhydrazone (CCCP) [46, 47]. Changing pH_*e*_ to either 8.15 or 8.45, we see that the magnitude of the PMF drops and then collapses to zero when we add CCCP. Intracellular pH rises with the alkaline shift and then starts to recover, but ΔpH is close to zero with CCCP, as predicted by the antiporter model. In Fig. S10 we experimentally confirm that sufficient amount of intracellular ATP is available to power any potential ATPases for ∼10 minutes after CCCP addition. The recovery of pH_*i*_ we see upon the alkaline shift at the lower magnitude PMF (Figs. 3C) agrees with previous measurements [48]. If we use the known range of the PMF during respiration on glucose to estimate Δ*ψ* (Fig. S11), Δ*ψ* stays approximately constant during the alkaline shifts, supporting a previously reported ∼ 10 mV change [48]. As expected in the antiporter model, Δ*ψ* is negligible in the absence of PMF (Fig. 3D).

### The optimal choice of antiporter depends on extracellular pH

We can extend our model to predict when cells should use each type of antiporter. For a given PMF and pH_*e*_, multiple different antiporters are in principle able to maintain the same steady state (Fig. 1D), and a cell could express, or upregulate the expression and/or activity of favoured antiporter [6, 30]. To identify preferred antiporters, we estimate each antiporter’s energetic cost.

We define this cost as the flux of protons that are neither directly involved in generating ATP during respiratory growth nor are actively pumped by F_1_F_*o*_ during fermentative growth. These protons enter the cell via leakage and the antiporters. They are costly either because they could have generated ATP during respiration or because they undermine F_1_F_*o*_’s export of protons during fermentation. We consider three antiporters, all with equivalents in *E. coli* (Fig. 1B), and write the cost as

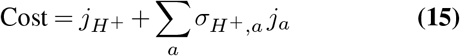

where as before the flux from leakage is 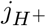 and from an antiporter *a* is *j*_*a*_.

The cost of maintaining a given PMF is minimised by using the antiporter with the smallest value of 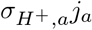, where Eq. (8) and Eq. (11) determine the value of *j*_*a*_ (*Supplementary Information*). For a minimal cost, cells should use each type of antiporter in a specific range of pH_*e*_ (Fig. 4A & B): the ClcA-like antiporter in a pH_*e*_ between approximately 2 and 5; the NhaB-like antiporter between 5 and 9; and the NhaA-like one between 9 and 12. Consistently, ClcA is important at acidic pH [49] and NhaA in alkaline pH [50, 51, 29], with its activity increasing once pH_*e*_ goes above 6.5 [52].

**Fig. 4.**
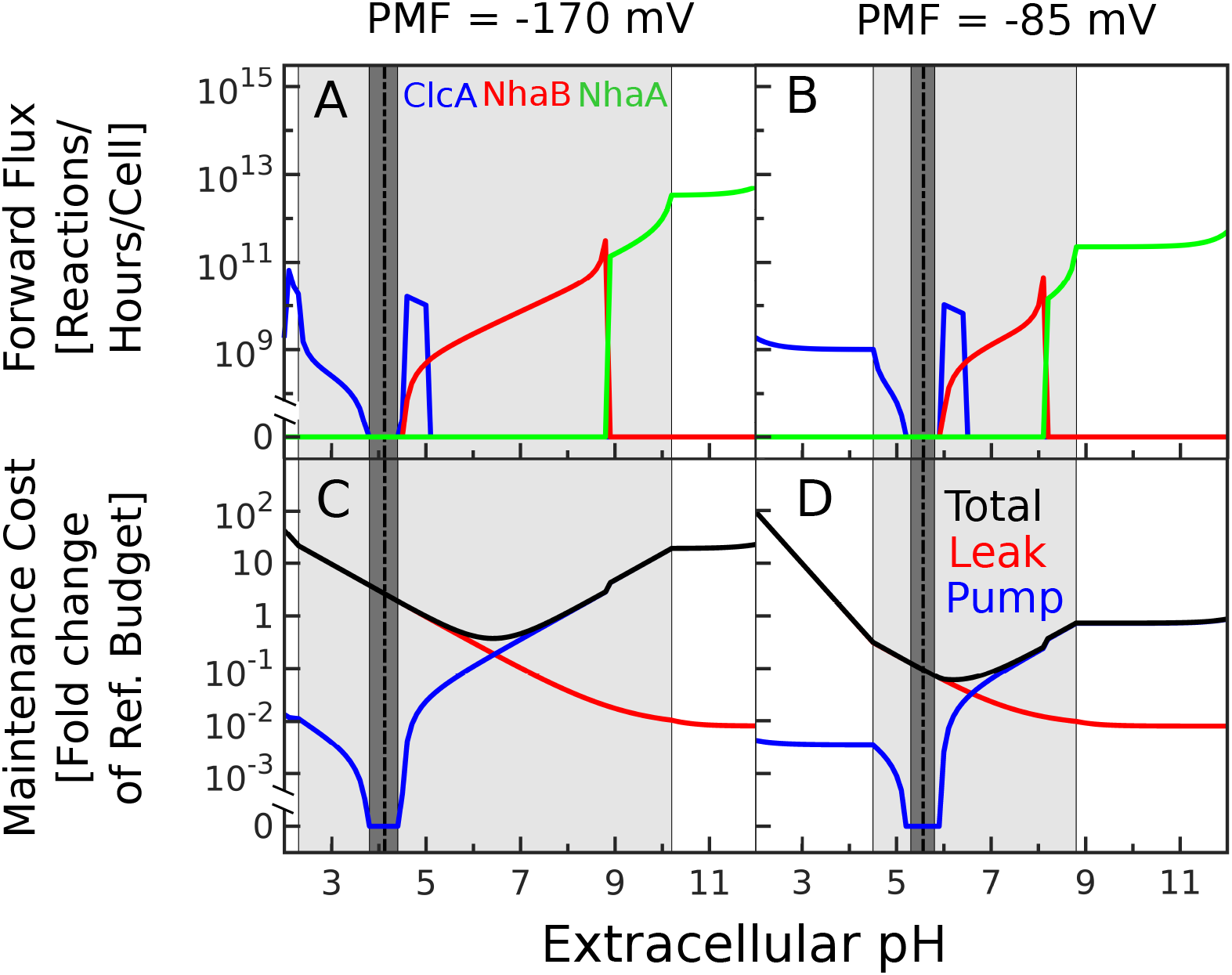
Maintaining pH_*i*_, the PMF, and osmotic pressure at minimal cost predicts a homeostatic strategy for *E. coli* across different pH_*e*_. We maintain pH_*i*_ as close as possible to 7, −10 ≤ *z*_*Y*_ ≤ 10, and set the osmotic pressure to 1 atm and the extracellular [CA]_0_ to 50 mM. The black dotted line marks the pH_*e*_ for which Δ*ψ* = 0; to the left, Δ*ψ >* 0; to the right, Δ*ψ <* 0. The dark grey indicates where the optimal solution has 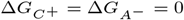 and so requires no antiporters. The light grey area indicates where pH_*i*_ can be maintained exactly at 7; to the left, pH_*i*_ *<* 7; to the right, pH_*i*_ *>* 7. (A,B) For two PMFs, we show the antiporter and its flux that minimises Eq. (15) for different pH_*e*_: a ClcA-like antiporter is in blue, a NhaB-like one in red, and a NhaA-like one in green. (C,D) For the same PMFs, we show the minimal cost (black) in protons per cell per hour relative to a reference budget, which we estimate for *E. coli* doubling every 60 min aerobically in minimal medium supplemented with glucose (SI). We also show the contributions to the cost of the antiporters’ activity (blue) and of leakage (red).

As pH_*e*_ changes so too does the dominant factor in the cost. For an alkaline pH_*e*_, the antiporters dominate the minimal cost; at acidic pH_*e*_, the leakage of protons dominates (Fig. 4C & D).

When minimising the cost, we also optimise the concentration and valency of the captive charges. For a small range of pH_*e*_, the cost is minimised by generating Δ*ψ* from the captive charges alone (Fig. 4), but this option is near to equilibrium and likely irrelevant physiologically. The optimal valency alters sign as pH_*e*_ changes (Fig. S13) and should be positive for sufficiently acidic pH_*e*_. This prediction is consistent with cells responding to acidic stress by generating positive charge: they convert some amino acids into more positively charged molecules and negatively charged glutamate into neutral *γ*-aminobutyric acid [53, 54, 55].

## Discussion

How bacteria maintain homeostasis has long been of interest. Here we gain new insights by noticing that protons and hydroxide ions contribute little to the membrane potential at a near neutral pH_*i*_. We identify that export of other ions is necessary. With no evidence for ATP-driven cation efflux in

*E. coli*, we predict that proton antiporters generate the cell’s out-of-equilibrium membrane potential. As a consequence, the PMF, which powers these antiporters, is necessary to maintain ΔpH and Δ*ψ*, and it determines how robustly cells can maintain their pH_*i*_ for different pH_*e*_. We confirm this prediction by demonstrating that collapsing the PMF depolarises the cell (Fig. 3) and that a lower magnitude PMF impairs *E. coli*’s maintenance of pH_*i*_ (Fig. 2).

Our results suggest that we may reconcile contradictory observations on the dynamics of pH_*i*_, [7, 8, 9] and [10, 11], if the cells in one experiment had a lower magnitude PMF, perhaps because of anoxic conditions.

Our prediction that proton:ion antiporters regulate Δ*ψ* suggests a shift of perspective away from electrogenicity, the net amount of charge transferred by an ion transporter, to the stoichiometric ratio of a given antiporter. If antiporters transport protons to keep pH_*i*_ neutral and Δ*ψ* fixed, as previously thought, the cell would lose control of the PMF as pH_*e*_ changes, which is not what we observe (Fig. 2D—F). Importantly, cells would need some other cation efflux pumps, apart from those for protons, to change ion concentrations to generate a physiological Δ*ψ*. In our perspective, even an electroneutral antiporter can generate Δ*ψ*, and ClcA, rather than exporting protons to raise pH_*i*_, exports chloride ions and imports protons at low extracellular pH (Fig. 1C & D). Alternatively, cells could actively maintain chloride motive force or actively import cations, such as by potassium ATPases [56], but we show that to keep the osmotic pressure in check the more likely strategy is to export anions (*Supporting Information*). Indeed, driven export of chloride occurs in the acidophile *Bacillus coagulans* [57].

This change of perspective applies too to alkalinophiles, which maintain a more negative Δ*ψ* compared to neutralophiles [1] and often have proton-sodium antiporters [58]. If those antiporters are responsible for generating the plasma membrane potential as we suspect, then we predict that their distinctive characteristic should be a high proton-to-sodium stoichiometric ratio.

Our work has caveats. It neither predicts the preferred pH_*i*_ nor addresses how cells recover this pH_*i*_ after a change in pH_*e*_, but rather we determine if a steady state with a neutral pH_*i*_ is possible. We therefore do not need to include how cells regulate the activity and expression of antiporters [6, 30] as they make this transition. Any direct regulation, which we do expect not only because our model does not predict the preferred pH_*i*_ but also because antiporters activity and expression is known to be pH_*i*_ regulated [6, 30], will be bound by the role of the antiporters on Δ*ψ* maintenance.

Similarly, we assume that the cell always has enough metabolic resources to actively export protons at a rate sufficient to maintain the PMF. The model is deliberately simple to be clear, focusing on one type of cation and one type of anion, both with a single charge. Multiple types of ions are of course present, but our approach remains qualitatively valid if cells regulate the membrane potential principally through one type of cation or anion whose active transport is mostly by one type of antiporter. We only implicitly model chemical reactions that modify pH_*i*_, such as the decarboxylation and carboxylation of amino acids and their transport [6, 53]. We capture these effects by changing the valency and concentration of captive molecules and so indubitably miss much [6]. Finally, our modelling does not address how cells move from one steady state to another because of a change in extracellular pH, but rather what steady states a particular antiporter supports.

Moving away from bacteria, proton-ion antiporters are widespread, and defects in these antiporters in mammalian cells are associated with disease [59, 60, 61]). In mammals, the members of the NHE gene family, to which *NhaA* belongs, and the CLC gene family, to which *ClcA* belongs, are also thought to help maintain cytoplasmic pH [62, 63]. Nevertheless, the multiple compartments of eukaryotic cells may mean that sustaining a PMF between the cytoplasm and extracellular space is not under such strong selection as for bacteria and archaea, constraining the activity of proton:ion antiporters less.

## Methods

### Strains and media

For simultaneous measurements of pH_*i*_ and the single-cell motor speed we use two strains of *E. coli*. The first is a derivative of MG1655 derivative (EK07 [34]) with sticky filaments and chromosomal expression of pHluorin. The second is the AB1157 strain [45] modified to carry the sticky flagellin encoding gene *fliC*^*sticky*^ [39, 34] and transformed with pkk223-3-pHluorin (M153R) plasmid [33]. FliC^*sticky*^ makes filaments hydrophobic so that they readily stick to polystyrene beads [39]. With the two strains, we confirmed that there are no significant strain-to-strain differences in the behaviour we observe. We used strain EK07 for pH_*e*_=5.5 experiments; strains EK07 and AB1157 for a pH_*e*_ of 5.7 and 6.0; strain AB1157 for a pH_*e*_ of 4, 4.3, 4.5, 4.7, 5, and 5.3; and strain EK07 for a pH_*e*_ of 8.15 and 8.45. MG1655 carrying pWR20-Q7* (Kanamycin resistance) and wild-type MG1655 was used for ATP measurements [64].

Bacteria were diluted (*×*10^−3^) from the frozen overnight culture (OD≈3) and grown in lysogeny broth (LB: 10 g tryptone, 5 g yeast extract, 10 g NaCl per 1 l) in conical flask at 37°C with shaking (220 rpm).

### Flagellar motor speed recording and analysis

Cells were harvested at OD= 2.0 to maximise the fraction that expressed the flagellar motor (Spectronic 200E Spectrophotometer, Thermo Scientific, UK). Cells were prepared for microscopy as before [65] using tunnel slides [66]. Experiments were performed in the motility buffer (BMB: aqueous solution of 6.2 mM K_2_HPO_4_, 3.8 mM KH_2_PO_4_, 67 mM NaCl, and 0.1 mM EDTA, pH 7.02) supplemented with 20 mM glucose.

*For aerobic-anaerobic shifts* (Fig. 2B & S5), cells were kept in glucose-supplemented BMB in the tunnel slide, where fresh BMB was flushed in just prior to the recording and the slide was sealed with CoverGrip™ Coverslip Sealant (Biotium, USA) to prevent oxygen diffusion. *For experiments presented in Fig. 2 C-F and S9*, bead rotation was recorded for 2 min in BMB with glucose in aerobic conditions, at which point the buffer was replaced with the pH-adjusted BMB. The slide was immediately sealed and recording continued for another 30 min. pH 5.5 buffer was adjusted with a mixture of organic acids (0.9 mM acetate, 3.8 mM lactate, 1.4 mM formate and 0.45 mM succinate), whereas the rest of the buffers were pH adjusted using lactic acid only. The choice of the organic acid influences only the dynamics of the drop in motor speed (such as in Fig. 2E), and not the steady speed. The recording of motor speed was uninterrupted for the duration of the experiment (32 min). For alkaline shifts (Fig. 3), we recorded bead rotation for 2 min before flushing BMB adjusted with NaOH to pH 8.15 or 8.45. The slide was not sealed to allow another flush of BMB supplemented with 100 μM CCCP at 17 min and recording was continued for another 10 min. *For experiments in Fig. S7* motility buffer was prepared to the required pH by adjusting the K_2_HPO_4_ and KH_2_PO_4_ ratio and if needed with KOH. 40 mM potassium benzoate and 40 mM methylamine hydrochloride (PBMH) was used to equilibrate pH_*e*_ and pH_*i*_ [9]. The experimental protocol was simiar to [34]. Briefly, tunnel slide was prepared in BMB of a given pH_*e*_, PBMH was added 1 min before adding a given concentration of butanol and 1 min after the beginning of the speed recording. Butanol was then washed out 1 to 2 min later. Finally, PBMH was washed out and the sequence was repeated for each recording. The motor speed in PBMH was compared with the speed during butanol flush.

Relative (*x, y*) bead coordinates recorded with a position-sensitive detector were converted to the flagellar motor rotational frequencies by applying a flat-top discrete Fourier transform with 1.6384 s window and 0.1 s step size [34, 41]. The resulting time series were median-filtered with the additional removal of 50 Hz AC frequency values and values below 10 Hz [34, 41].

### Intracellular pH measurements

The intracellular pH of the bacteria was measured with ratiometric pHluorin [67, 38] using an established imaging protocol and sensor calibration [68]. Images were taken every 90 s with 50 ms exposure time.

Fluorescence images of pHluorin were analysed as before [34, 38]. An Otsu threshold followed by binary erosion and binary dilation was applied to both 475 nm and 395 nm excitation images, and the mean intensity was calculated for each cell in both channels. The ratio of these values with the background subtracted was later converted to pH units using a calibration curve generated following [38].

### Intracellular ATP measurements

Cells were harvested at OD=2, washed 3 times by centrifugation at 8000g for 2 min, and resuspended in BMB with 20 mM glucose, adjusted to pH 8.13-8.2 with KOH, and at final OD=1. Cells were transferred to the Black Nunc 96-Well Plate (Flat Bottom), where 100 μM of CCCP was added with the multichannel pipette, simultaneously to all but one well, which served as a control. The measurements were performed using Biotek Synergy H1 plate reader, in 20, 60 s or 5 min intervals post CCCP addition, and on each well once (to avoid reported photoactivation of Queen7 μM* [64]). Parallel measurements were performed with Wild type MG1655 to correct for background fluorescence by subtracting it from Queen7 μM* signal. To minimise photoactivation of neighbouring wells in the plate reader measurements were repeated 4 times, and taken in a different well order. Cells were excited with 405 and 488 nm wavelengths, and emission was taken at 528 nm. The 405/488 ratios were converted to the ATP concentrations according to the calibration curve given. All experiments were performed at room temperature.

*Queen7* μM*\* sensor calibration* was performed as before [64]. Cells were grown and prepared as before to Queen buffer (50 mM HEPES, 200 mM KCl, 1 mM MgCl_2_, 0.05% Triton X-100, protease inhibitor cocktail (Sigma-Aldrich)) adjusted to pH 8.13. 100 μg/ml of lysozyme were added to the cells and incubated at room temperature for 15 min. The culture was then frozen to -70°C and thawed twice to weaken the cell wall. Cells were later broken down by sonicating on ice at 4°C with Qsonica Q700 at amplitude 50, 4 pulses of 30 s with 30 s interval (∼1000 J total energy input). Cell debris were spun down at 13000g for 1 min and the supernatant was filtered with 0.22 μm syringe filter. Lysate fluorescence, containing Queen7 μM* and a known concentration of ATP (Adenosine 5’-triphosphate magnesium salt, Sigma-Aldrich), was measured at 405 and 488 nm excitation and 528 nm emission wavelengths. MG1655 lysate was used for background fluorescence measurements. The calibration curve was fitted with the sigmoid 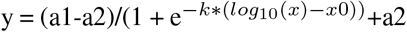, Fig. S10B.

### Computational results

The computational algorithms, including useful heuristic approximations we developed, are described in the *Supplementary Information*. To numerically solve the steady-state equations we used the MATLAB function `vpasolve`.

### Availability of data

Our experimental data is available at https://datashare.is.ed.ac.uk/handle/10283/2058.

## ACKNOWLEDGEMENTS

We would like to thank all members of the Pilizota and Swain labs as well as Sebastian Jaramillo-Riveri, Xavier Zaoui, Vincent Danos, Joshua Shaevitz, Ariel Amir, and Calin Guet for their support and useful discussions. We would like to thank Joanne Slonezewski and Linda Kenny for providing and helping us with reproducing their previously published data in Fig. 1A. TP, GT and EK were supported by the Human Frontier Science Program grant (RGP0041/2015), GT is supported by the Darwin Trust of the University of Edinburgh, and PSS by the BBSRC.

## Notes

### Competing Interest Statement

The authors have declared no competing interest.

### Summary of Updates

The revision includes additional data and figures. The text has also been rewritten for clarity.

