## Supplementary material for "The proton motive force determines *Escherichia coli*’s robustness to extracellular pH": SI text and figures

### Supplementary text

**Cell shape.** To calculate a cell's surface area and volume, we assume that the cell is a spherocylinder with a length-to-width ratio of 3:1, determined from our microscopy images [1] and consistent with [2], and write:

$$S = \pi w l \tag{S1}$$

$$V = \pi \left(\frac{w}{2}\right)^2 \left(l - \frac{w}{3}\right) \tag{S2}$$

where $l$, $w$, and $V$ are the cell's length, width, and volume. When calculating osmotic pressure, we assume that cellular volume stays constant and ignore cell wall mechanical properties for simplicity.

**Describing the movement of ions.** To describe the movement of ions across the plasma membrane, we use Eq. 5, which is

$$\frac{d[x]_i}{dt} = -\frac{d[x]_e}{dt} = j_x + \sum_a \sigma_{x,a}\, j_a \tag{S3}$$

where $j_x$ is the flux of ions due to leakage and $j_a$ is the flux generated by pumping reaction $a$ with a stoichiometric coefficient of $\sigma_{x,a}$ (Table S1).

We assume that all transport and leakage reactions are reversible and that the corresponding flux can be decomposed into the rate of import into the cell, and the rate of export. For a transporter this is $j_a^+$ and $j_a^-$:

$$j_a = j_a^+ - j_a^- = j_a^+ \left(1 - \frac{j_a^-}{j_a^+}\right) \tag{S4}$$

where $j_a^+ \geq 0$ and $j_a^- \geq 0$. Similar would hold true for leakage.

The Gibbs free energy of this reaction, $\Delta G_a$ (Table S1), can be written as [3, 4]

$$\Delta G_a \equiv \frac{RT}{F} \ln\left(\frac{j_a^-}{j_a^+}\right), \tag{S5}$$

measuring $\Delta G_a$ in volts, using the definition of $\Delta G_a$ and writing $\Delta G_0 = -\log(K_{\text{eq}})/RT$ with detailed balance implying that the equilibrium constant $K_{\text{eq}}$ is the ratio of the forward to backward rate constants. Eq. S4 then becomes

$$j_a = j_a^+ \left(1 - e^{\frac{F}{RT}\Delta G_a}\right) \tag{S6}$$

where $j_a^+$ is a function determined by the specifics of the transport [5].

***Leakage reactions.*** For the leakage flux of ion $x$, we again use Eq. 8 in the main text, where j$^+$ is given as:

$$j_x^+ = \frac{S}{V} P_x x_e f_b(\Delta\psi) \tag{S7}$$

where $P_x$ the membrane's permeability to $x$, and the functional form of $f_b(\Delta\psi)$ changes with the choice of approximation [6]. To describe the flux of ions from leakage we choose the trapezoidal energy barrier approximation [6] for all ions, which gives $f_b$ as

$$f_b(z_x\Delta\psi) = \frac{\frac{F}{RT} b z_x \Delta\psi \cdot \mathrm{e}^{-\frac{F}{RT} z_x \Delta\psi/2}}{\mathrm{e}^{b\frac{F}{RT} z_x\Delta\psi/2} - \mathrm{e}^{-b\frac{F}{RT} z_x\Delta\psi/2}}. \tag{S8}$$

Here $b$ is the fractional width of the trapezoid, $0 \leq b \leq 1$, which specifies how the electrostatic potential varies within the membrane [6].
Finally, inserting Eq. S7 and Eq. S8 into the Eq. 8 in the main text we get:

$$j_x = \frac{S}{V} P_x [x]_e f_b(z_x\Delta\psi)\left(1 - \mathrm{e}^{\frac{F}{RT}\Delta G_x}\right) \tag{S9}$$

When $b = 0$, Eq. S9 is Erying's single barrier model:

$$j_x(b=0) = \frac{S}{V} P_x [x]_e e^{-\frac{F}{RT} z_x \Delta\psi/2}\left(1 - e^{\frac{F}{RT}\Delta G_x}\right) \tag{S10}$$

when $b = 1$, Eq. S9 is the Goldman–Hodgkin–Katz flux equation [6]. We use latter to compute the cost Erying's equation above.

***The dynamics of protons and the dissociation of water.*** In our model, protons are the only ions whose pumping is powered by the cell's metabolism. Their electrochemical potential then powers all antiporters, making their dynamics the most complex. Using Table S1, we have

$$\frac{d[\mathrm{H_3O^+}]_i}{dt} = \underbrace{j_{\mathrm{H^+}}}_{leakage} \underbrace{-10 j_{\mathrm{R}} + 3.3 j_{\mathrm{F1fo}} + 3 j_{\mathrm{NhaB}} + 2 j_{\mathrm{NhaA}} + j_{\mathrm{ClcA}}}_{pumping} + f_w [\mathrm{H_2O}]_i^2 - b_w [\mathrm{H_3O^+}]_i [\mathrm{OH^-}]_i \tag{S11}$$

where $f_w$ and $b_w$ are rate of association of $H_3O^+$ and $OH_-$ and dissociation rate of $H_2O$, respectively.
Hydroxide ions, on the other hand only leak according to

$$\frac{d[\mathrm{OH^-}]_i}{dt} = \underbrace{j_{\mathrm{OH^-}}}_{leakage} + f_w [\mathrm{H_2O}]_i^2 - b_w [\mathrm{H_3O^+}]_i [\mathrm{OH^-}]_i \tag{S12}$$

In practice, we use Eq. S11 only at steady state and consequently assume that the hydroxide ions are also at steady state:

$$j_{\mathrm{OH^-}} + f_w [\mathrm{H_2O}]_i^2 - b_w [\mathrm{H_3O^+}]_i [\mathrm{OH^-}]_i = 0 \tag{S13}$$

so that the terms from the ionisation of water in Eq. S11 can be replaced:

$$\frac{d[\mathrm{H_3O^+}]_i}{dt} = j_{\mathrm{H^+}} - 10 j_{\mathrm{R}} + 3.3 j_{\mathrm{F1fo}} + 3 j_{\mathrm{NhaB}} + 2 j_{\mathrm{NhaA}} + j_{\mathrm{ClcA}} - j_{\mathrm{OH^-}} \tag{S14}$$

To compute $f_w$ in Eq. S13, we assume that the rates $f_w$ and $b_w$ are the same intracellularly and extracellularly. If $[\mathrm{H_2O}]_e = 1000/18$ M and $b_w$, the rate of association of hydronium and hydroxide ions, is diffusion-limited, so that $b_w = 10^{10}$ M$^{-1}$ s$^{-1}$ [7], then $f_w = K_w b_w / [\mathrm{H_2O}]_e^2$ or approximately 3.2 $\times 10^{-8}$ M$^{-1}$ s$^{-1}$, assuming $K_w = 10^{-14}$ M$^2$.

***Bounds on the PMF.*** To determine physiological limits on the PMF we refer to the literature and again take a minimal approach. We define the respiratory regime by the coupling of the redox reaction $\mathrm{R^\star} \rightleftharpoons \mathrm{R}$, where $\mathrm{R^\star}$ is typically NADH, to the export of protons:

$$\mathrm{R^\star} + \sigma_{H^+,R}\mathrm{H}_i^+ \rightleftharpoons \mathrm{R} + \sigma_{H^+,R}\mathrm{H}_e. \tag{S15}$$

with $\sigma_{H^+,R}$ being the stoichiometric coefficient determining the number of protons moved across the membrane for each $\mathrm{R^\star}$. Writing $\Delta G_{\mathrm{NADH}}$ as the redox reaction's potential, then

$$\Delta G_R = \Delta G_{\mathrm{NADH}} - \sigma_{H^+,R}\Delta G_{H^+} < 0 \tag{S16}$$

if protons are to be exported. These protons are used by the $F_1F_o$ ATP synthase to generate ATP [8]:

$$\mathrm{ADP} + \sigma_{\mathrm{H},F_1F_o}\mathrm{H}_e \rightleftharpoons \mathrm{ATP} + \sigma_{\mathrm{H},F_1F_o}\mathrm{H}_i, \tag{S17}$$

and for the flux of ATP to be positive, we require:

$$\Delta G_{F_1F_o} = \sigma_{H+,F_1F_o}\Delta G_{H+} - \Delta G_{\mathrm{ATP}} < 0. \tag{S18}$$

Together Eq. S16 and Eq. S18 bound the magnitude of the PMF in the respiratory regime:

$$\frac{\Delta G_{\mathrm{NADH}}}{\sigma_{H+,R}} < \Delta G_{H+} < \frac{\Delta G_{\mathrm{ATP}}}{\sigma_{H+,F_1F_o}} \tag{S19}$$

To estimate these limits, we use $\Delta G_{\mathrm{ATP}} \simeq$ -560 mV for exponentially growing cells respiring on glucose [9] and $\sigma_{H+,F_1F_o} =$ 3.3 [10] (see also ***E. coli* energy budget** below). For $\Delta G_R$, we use -2290 mV from the Gibbs free energy of the half-reactions [11] and with 10 protons moved per NADH oxidised [12]. We thus find that

$$-229 \text{ mV } < \Delta G_{H+} < -170 \text{ mV }. \tag{S20}$$

The fermentative regime is coarse grained by representing it with $F_1F_o$ hydrolysing ATP to export protons, which requires $\sigma_{H+,F_1F_o}\Delta G_{H+} > \Delta G_{\mathrm{ATP}}$ or

$$\Delta G_{H+} > -170 \text{ mV }. \tag{S21}$$

A caveat is that the free energy of ATP hydrolysis and the stoichiometric ratio $\sigma_{H+,F_1F_o}$ may change in different extracellular environments [13, 14], which would modify the value of the PMF that separates the respiratory and fermentative regimes. For example, in Fig. 3D in the main text, although we obtained data in the presence of oxygen, we estimate the PMF at which we observe $\mathrm{pH}_i$ recovery to be in the fermentative regime. Thus, the free energy and stoichiometric ratio of ATP hydrolysis or of NADH reduction could have changed at high $\mathrm{pH}_e$. Alternatively, cells in Fig. 3D in the main text may have switched to fermentation in alkaline stress.

**Extracellular environment.** We start with a solution of water with a pH of 7. To change $\mathrm{pH}_e$, we add either acid $[\mathrm{AH}]_0$ or base $[\mathrm{COH}]_0$. To change the concentration of extracellular salt, we add $[\mathrm{CA}]_0$. Assuming equilibrium, we use the known dissociation and associations constants, conservation of mass, and the charge neutrality of the solution to define fully the extracellular environment.

Water, hydroxide ions, and hydronium ions obey

$$2\mathrm{H_2O} \rightleftharpoons \mathrm{H_3O^+} + \mathrm{OH^-} \tag{S22}$$

and defining the equilibrium constant of this reaction as $K_c$, we have that

$$K_c = \frac{[\mathrm{H_30^+}]_e[\mathrm{OH^-}]_e}{[\mathrm{H_2O}]_e^2}. \tag{S23}$$

Taking logarithms and writing $K_w = K_c[\mathrm{H_2O}]_e^2$, which is sometimes called the ionic product of water, then

$$\mathrm{pK}_w = \mathrm{pH}_e + \mathrm{pOH}_e \tag{S24}$$

where we use the prefix p to denote $-\log_{10}$. If we assume that $[\mathrm{H_2O}]_e$ is the molar concentration of water (approximately 1000/18 M) $\mathrm{pK}_w$ is approximately 14 [7].

The acid dissociates

$$\mathrm{AH} + \mathrm{H_2O} \rightleftharpoons \mathrm{A^-} + \mathrm{H_3O^+} \tag{S25}$$

with an equilibrium dissociation constant of $K_c$

$$K_c = \frac{[\mathrm{H_30^+}]_e[\mathrm{A^-}]_e}{[\mathrm{H_2O}]_e[\mathrm{AH}]_e}. \tag{S26}$$

and so a $\mathrm{pKa} = -\log_{10}(K_c[\mathrm{H_2O}])$ of

$$\mathrm{pKa} = -\log_{10}\left(\frac{[\mathrm{A^-}]_e[\mathrm{H_3O^+}]_e}{[\mathrm{AH}]_e}\right). \tag{S27}$$

Similarly, the base dissociates too

$$\mathrm{COH} \rightleftharpoons \mathrm{C}^{+} + \mathrm{OH}^{-} \tag{S28}$$

with a pKb of

$$\mathrm{pKb} = -\log_{10}\left(\frac{[\mathrm{C}^{+}]_e[\mathrm{OH}^{-}]_e}{[\mathrm{COH}]_e}\right). \tag{S29}$$

From conservation of mass, and assuming that the salt fully dissociates once added to the external environment

$$\mathrm{CA} \longrightarrow \mathrm{A}^{-} + \mathrm{C}^{+} \tag{S30}$$

so that $[\mathrm{CA}]_e \approx 0$, we have

$$\begin{aligned} [\mathrm{COH}]_0 + [\mathrm{CA}]_0 &= [\mathrm{COH}]_e + [\mathrm{CA}]_e + [\mathrm{C}^{+}]_e \\ &\simeq [\mathrm{COH}]_e + [\mathrm{C}^{+}]_e \end{aligned} \tag{S31}$$

$$\begin{aligned} [\mathrm{AH}]_0 + [\mathrm{CA}]_0 &= [\mathrm{AH}]_e + [\mathrm{CA}]_e + [\mathrm{A}^{-}]_e \\ &\simeq [\mathrm{AH}]_e + [\mathrm{A}^{-}]_e. \end{aligned} \tag{S32}$$

Lastly, charge neutrality implies that

$$[\mathrm{H_3O}^{+}]_e - [\mathrm{OH}^{-}]_e + [\mathrm{C}^{+}]_e - [\mathrm{A}^{-}]_e = 0. \tag{S33}$$

Given $\mathrm{pH}_e$, $[\mathrm{AH}]_0$, $[\mathrm{COH}]_0$ and $[\mathrm{CA}]_0$, we then use Eqs. S24 to S33 to find the remaining concentrations. First, from Eq. S24,

$$[\mathrm{OH}^{-}]_e = 10^{\mathrm{pH}_e - \mathrm{pK}_w} \tag{S34}$$

and, using Eq. S29 and Eq. S34, Eq. S31 becomes

$$[\mathrm{COH}]_0 + [\mathrm{CA}]_0 = \left(1 + 10^{\mathrm{pKb}+\mathrm{pHe}-\mathrm{pK}_w}\right)[\mathrm{C}^{+}]_e \tag{S35}$$

and, using Eq. S27 and definition of $\mathrm{pH}_e$, Eq. S32 becomes

$$[\mathrm{AH}]_0 + [\mathrm{CA}]_0 = \left(1 + 10^{\mathrm{pKa}-\mathrm{pHe}}\right)[\mathrm{A}^{-}]_e. \tag{S36}$$

Second, if $\mathrm{pH}_e$ is acidic, we do not add base so that $[\mathrm{COH}]_0 = 0$. Then, we can express $[C^{+}]_e$ from Eq. S35 and using Eq. S33 we get

$$[\mathrm{A}^{-}]_e = \frac{[\mathrm{CA}]_0}{1 + 10^{\mathrm{pKb}+\mathrm{pHe}-\mathrm{pK}_w}} + 10^{-\mathrm{pH}_e} - 10^{\mathrm{pH}_e - \mathrm{pK}_w}. \tag{S37}$$

Third, if $\mathrm{pH}_e$ is alkaline, we do not add acid so that $[\mathrm{AH}]_0 = 0$. Then, we can express $[A^{-}]_e$ from Eq. S36 and using Eq. S33 we get

$$[\mathrm{C}^{+}]_e = \frac{[\mathrm{CA}]_0}{1 + 10^{\mathrm{pKa}-\mathrm{pH}_e}} - 10^{-\mathrm{pH}_e} + 10^{\mathrm{pH}_e - \mathrm{pK}_w}. \tag{S38}$$

By assuming the salt fully dissociates, we have therefore completely specified the extracellular environment by Eq. S34, Eq. S37, and Eq. S38 given only $[\mathrm{CA}]_0$ and $\mathrm{pH}_e$.

**Model reactions.** Summary of all the model's reactions

| Flux | Reaction | Free Energy Change | Refs. (for stoichiometries) |
|---|---|---|---|
| $j_w$ | $2H_2O_i \rightleftharpoons H_3O_i^+ + OH_i^-$ | NA | NA |
| $j_{H^+}$ | $H_e^+ \rightleftharpoons H_i^+$ | $\Delta G_{H^+}$ | NA |
| $j_{OH^-}$ | $OH_e^- \rightleftharpoons OH_i^-$ | $\Delta G_{OH^-}$ | NA |
| $j_{C^+}$ | $C_e^+ \rightleftharpoons C_i^+$ | $\Delta G_{C^+}$ | NA |
| $j_{A^-}$ | $A_e^- \rightleftharpoons A_i^-$ | $\Delta G_{A^-}$ | NA |
| $j_R$ | $R^\star + 10\ H_i^+ \rightleftharpoons R + 10\ H_e^+$ | $\Delta G_R = \Delta G_{NADH} - 10\Delta G_{H^+}$ | [15] |
| $j_{F1F_o}$ | $ATP + 3.3\ H_e^+ \rightleftharpoons ADP + 3.3\ H_i^+$ | $\Delta G_{F1F_o} = \Delta G_{ATP} + 3.3\Delta G_{H^+}$ | [10] |
| $j_{NhaB}$ | $3\ H_e^+ + 2C_i^+ \rightleftharpoons 3\ H_i^+ + 2\ C_e^+$ | $\Delta G_{NhaB} = 3\Delta G_{H^+} - 2\Delta G_{C^+}$ | [16] |
| $j_{NhaA}$ | $2\ H_e^+ + C_i^+ \rightleftharpoons 2\ H_i^+ + C_e^+$ | $\Delta G_{NhaA} = 2\Delta G_{H^+} - \Delta G_{C^+}$ | [17, 18] |
| $j_{ClcA}$ | $H_e^+ + 2A_i^- \rightleftharpoons H_i^+ + 2\ A_e^-$ | $\Delta G_{ClcA} = \Delta G_{H^+} - 2\Delta G_{A^-}$ | [19] |

**Table S1.** Summary of all the reactions we consider in the model.

**A mathematical model of the membrane potential.** To solve Eq. 2 in the main text, we first rewrite it as

$$\Delta\psi = \frac{FV}{SC_m}\left(z_Y[\mathrm{Y}]_i + [\mathrm{H}^+]_i - [\mathrm{OH}^-]_i + [\mathrm{C}^+]_i - [\mathrm{A}^-]_i\right) \tag{S39}$$

where $Y_i$ are the captive molecules with valency $z_Y$. Next we express $[\mathrm{C}^+]_i$ and $[\mathrm{A}^-]_i$ as a function of $[\mathrm{C}^+]_e$ and $[\mathrm{A}^+]_e$ using the definition of $\Delta G$ :

$$\Delta\psi = \frac{FV}{SC_m}\left(z_Y[\mathrm{Y}]_i + [\mathrm{H}^+]_i - [\mathrm{OH}^-]_i + [\mathrm{C}^+]_e \cdot \mathrm{e}^{\frac{F}{RT}(\Delta G_{C^+} - \Delta\psi)} - [\mathrm{A}^-]_e \cdot \mathrm{e}^{\frac{F}{RT}(\Delta G_{A^-} + \Delta\psi)}\right). \tag{S40}$$

As discussed in the main text, we disregard the negligible contribution from $[\mathrm{H}^+]_i$ and $[\mathrm{OH}^-]_i$. Given our assumptions on *E. coli*'s shape, the surface to volume ratio is $\sim 4.25 \times 10^6\ \mathrm{m}^{-1}$, and using the previously reported $C_m \sim 6.5 \times 10^{-3}\ \mathrm{F\ m}^{-1}$ [20], we find that $\frac{FV}{SC_m} \sim 3.5$ V per mM.

With the above assumptions, $\Delta\psi$ becomes

$$\Delta\psi = 3.5 \cdot \left(z_Y[\mathrm{Y}]_i + [\mathrm{C}^+]_e \cdot \mathrm{e}^{\frac{F}{RT}(\Delta G_{C^+} - \Delta\psi)} - [\mathrm{A}^-]_e \cdot \mathrm{e}^{\frac{F}{RT}(\Delta G_{A^-} + \Delta\psi)}\right). \tag{S41}$$

Dividing Eq. S41 by $[\mathrm{Ion}]_e$, the total concentration of extracellular ions, we express Eq. S41 compactly as

$$\begin{aligned} 3.5 \cdot \frac{\Delta\psi}{[\mathrm{Ion}]_e} &= \alpha_Y + \alpha_{C^+} \cdot \mathrm{e}^{\frac{F}{RT}(\Delta G_{C^+} - \Delta\psi)} - \alpha_{A^-} \cdot \mathrm{e}^{\frac{F}{RT}(\Delta G_{A^-} + \Delta\psi)} \\ &= \alpha_Y + \sum_x z_x \alpha_x \cdot \mathrm{e}^{\frac{F}{RT}(\Delta G_x - z_x\Delta\psi)} \end{aligned} \tag{S42}$$

where $\alpha_x = [x_e]/[\mathrm{Ion}]_e$ is the fraction of either cation or anion in the environment, with $\sum_x \alpha_x = 1$, and $\alpha_Y = z_Y[\mathrm{Y}]_i/[\mathrm{Ion}]_e$ and with the total extracellular concentration of ions being

$$[\mathrm{Ion}]_e = [\mathrm{C}^+]_e + [\mathrm{A}^-]_e + [\mathrm{OH}^-]_e + [\mathrm{H}^+]_e. \tag{S43}$$

If pHe is acidic, then Eqs. S33, S34 and Eq. S37 imply that Eq. S43 becomes

$$\begin{aligned} [\mathrm{Ion}]_e &= 2\left([\mathrm{A}^-]_e + [\mathrm{OH}^-]_e\right) \\ &= 2\left(\frac{[\mathrm{CA}]_0}{1 + 10^{\mathrm{pKb} + \mathrm{pH}_e - \mathrm{pK}_w}} + 10^{-\mathrm{pH}_e}\right). \end{aligned} \tag{S44}$$

If $\mathrm{pH}_e$ is alkaline, then the definition of pH, and Eqs. S33 and Eq. S38 imply that Eq. S43 becomes

$$[\mathrm{Ion}]_e = 2\left([\mathrm{C}^+]_e + [\mathrm{H}^+]_e\right)$$
$$= 2\left(\frac{[\mathrm{CA}]_0}{1+10^{\mathrm{pKa}-\mathrm{pH}_e}} + 10^{\mathrm{pH}_e-\mathrm{pK}_w}\right). \quad \textbf{(S45)}$$

For sufficiently large $[\mathrm{CA}]_0$, we approximately solve Eq. S42 for $\Delta\psi$ both by setting the left hand side of Eq. S42, which is small, to zero and by using $\alpha_{C^+} = \alpha_{A^-} \approx 1/2$.
This approximation:

$$\alpha_Y = \frac{1}{2}\cdot\left(\mathrm{e}^{\frac{F}{RT}(\Delta G_{A_-}+\Delta\psi)} - \mathrm{e}^{\frac{F}{RT}(\Delta G_{C_+}-\Delta\psi)}\right) \quad \textbf{(S46)}$$

is robust for $\mathrm{pH}_e$ and $\mathrm{pH}_i$ in [4, 10] (Fig. S13) and we use it in Fig. 3.

$$\Delta\psi \simeq \frac{FV}{SC_m}\left(z_Y[\mathrm{Y}]_i + [\mathrm{C}^+]_e\cdot\mathrm{e}^{-\frac{F}{RT}\Delta\psi} - [\mathrm{A}^-]_e\cdot\mathrm{e}^{\frac{F}{RT}\Delta\psi}\right) \quad \textbf{(S47)}$$

whose solution must hold both at and away from equilibrium. Therefore the membrane potential can never deviate substantially from the equilibrium potential if both the intracellular concentrations of protons and hydroxide ions remain negligible and there is no active transport of ions.

**Antiporters with different stochiometries generate steady states with $\mathrm{pH}_i$=7 for different ranges of PMF and $\mathrm{pH}_e$.** To understand why, we consider an NhaA-like antiporter when $z_Y[Y]_i = 0$ in Eq. 10 in the main text and $\mathrm{pH}_e$ is 5.5 (Fig. 1D in the main text – black line). The corresponding $\Delta$pH is 1.5 and contributes -90 mV to the PMF. The antiporter exchanges cations ($x = C^+$) and $\Delta G_{A^-} = 0$. Consequently, $[A_i]$ passively follows the membrane potential with $[A_i] = [A_e]\exp(\frac{F\Delta\psi}{RT})$, increasing if $\Delta\psi$ becomes more positive, and vice versa.

For this NhaA-like antiporter, steady states become impossible for less negative PMFs (Fig. 1D in the main text), which, from Eq. 2 in the main text, require a more positive $\Delta\psi$ because $\Delta$pH is fixed. A more positive $\Delta\psi$ increases $[A_i]$, and so, from Eq' 9 in the main text, can only be generated if $[C_i]$ increases. This increase in $[C_i]$ and the more positive $\Delta\psi$ raises $\Delta G_{C^+}$ via Eq. 1 in the main text until the upper bound of Eq. 8 in the main text is reached, and cations begin to leak out of, not into, the cell. If $z_Y[Y]_i$ is negative rather than zero, then for a given, sufficiently positive PMF, a given $\mathrm{pH}_e$ and a neutral $\mathrm{pH}_i$, $[C_i]$ must be higher. These higher concentrations favour cations leaking from the cell and so lower the $\Delta\psi$ and hence the PMF at which the upper bound in Eq. 8 in the main text is reached (Fig. 1D top in the main text and S3).

Steady states also become impossible for sufficiently negative PMFs for an NhaA-like antiporter (Fig. 1D in the main text). Such PMFs require a more negative $\Delta\psi$ because $\Delta$pH is fixed. This $\Delta\psi$ decreases $[A_i]$ and so can only be generated if $[C_i]$ decreases too. The decrease in $[C_i]$ and the more negative $\Delta\psi$ lowers $\Delta G_{C^+}$. Exporting cations becomes harder until the lower bound of Eq. 8 in the main text is reached. The export of cations then stops, preventing a steady state. Although a more negative PMF does increase the free energy available to drive antiport, the corresponding decrease in $[C_i]$ dominates, and the overall free-energy change eventually becomes zero. If $z_Y[Y]_i$ is positive rather than zero, $[C_i]$ must be lower for a given PMF and $\mathrm{pH}_e$. These lower concentrations make exporting cations harder, and so the point where the PMF is decreased sufficiently to stop export through the corresponding decrease in $[C_i]$ occurs at a higher PMF (Fig. 1D bottom in the main text and S3).

**Energy budget of *E. coli*.** To calculate the reference energy budget of *E. coli* (Fig. 4D), we used that *E. coli* growing at 0.89 $\mathrm{h}^{-1}$ on glucose consumes 19.8 $\mathrm{mMh}^{-1}$ of oxygen per gram of dry weight [21]. When growing at 1 $\mathrm{h}^{-1}$, the dry weight of *E. coli* is 258 $\times$ 10 $^{-15}$ g per cell [22]. Assuming that the oxygen consumption and the mass per cell are comparable between the two experiments, we have that $19.8\times10^{-3}\times6\times10^{23}\times258\times10^{-15} = 3.08\times10^{9}$ $\mathrm{O}_2$ consumed per cell per hour. With a stoichiometry of the respiratory chain of $\sigma_{\mathrm{H},R} = 10$ per $1/2$ $\mathrm{O}_2$ [15], this consumption rate corresponds to a 'budget' of $61.6\times10^{10}$ protons per cell per hour.

**Modelling the periplasm.** We have considered a single barrier separating the cell's interior from its environment, but *E. coli* has an inner and an outer membrane [23]. The ionic pumps are all in the inner membrane [24, 25], and the outer membrane contains porins, which allow ions to move freely between the periplasm and the extracellular space [26, 27]. These ions should equilibrate so that the periplasmic and extracellular concentrations follow a Nernst equation. Setting Eq. 1 to zero, we obtain:

$$[x]_p = [x]_e\cdot\mathrm{e}^{-\frac{z_xF}{RT}\Delta\psi_p} \quad \textbf{(S48)}$$

where $[x]_p$ is the periplasmic concentration of an ion $x$ and $\Delta\psi_p$ is the equilibrium potential across the outer membrane, which will only be non-zero if the periplasm contains captive charged molecules.

From Eq. 1, 16 and Eq. S48, we find:

$$
\begin{aligned}
[x]_i &= [x]_p \cdot \mathrm{e}^{\frac{z_x F}{RT}(\Delta G_x - z_x \Delta\psi)} \\
&= [x]_e \cdot \mathrm{e}^{\frac{z_x F}{RT}(\Delta G_x - z_x \Delta\psi_p - z_x \Delta\psi)}
\end{aligned} \tag{S49}
$$

Eq. S49 shows that $\Delta\psi_p$ offsets the extracellular concentrations 'felt' by at the inner membrane, so that the ΔpH between the cytoplasm and the periplasm differs from the ΔpH between the cytoplasm and the extracellular environment (Fig. S14). A $\Delta\psi_p$ of -30 mV [28] increases a cationic and decreases an anionic concentration by $\sim$ 3.3 times.

Using Eq. S49, Eq. S41 becomes

$$
\Delta\psi = \frac{FV}{SC_m}\left[z_Y[Y]_i + \sum_{x \in \text{small ion}} [x]_e \cdot \mathrm{e}^{\frac{F}{RT}(\Delta G_x - z_x\Delta\psi - z_x\Delta\psi_p)}\right]. \tag{S50}
$$

### Computational experiments.

***Constants.*** All constants are specified in Table S2.

| Name | Symbol | Units | Value | Ref. |
|---|---|---|---|---|
| Faraday constant | $F$ | C/mole | 96485 | |
| Gas constant | $R$ | J/mole/K | 8.31 | |
| Temperature | $T$ | K | 298 | |
| Dissociation constant HCl | $pKa_{AH}$ | Dimensionless | -6.3 | |
| Dissociation constant NaOH | $pKb_{COH}$ | Dimensionless | -0.56 | |
| Dissociation constant $H_2O$ | $pKa_w$ | Dimensionless | 14 | |
| Association rate of $H_3O^+$ and $OH^-$ | $f_w$ | $M^{-1}s^{-1}$ | $10^9$ | |
| Dissociation rate of $H_2O$ | $b_w$ | $M^{-1}s^{-1}$ | $3{\cdot}10^{-9}$ | |
| Membrane capacitance | $C_m$ | C $m^{-2}$ | $6.5\times10^{-3}$ | [20] |
| Cell width | $w$ | m | $1.07\times10^{-6}$ | [2, 1] |
| Cell length | $l$ | m | $2.95\times10^{-6}$ | [2, 1] |
| Gibbs free energy of NADH reduction | $\Delta G_{\mathrm{NADH}}$ | V | -2.290 | [11] |
| Gibbs free energy of ATP hydrolysis | $\Delta G_{\mathrm{ATP}}$ | V | -0.560 | [11] |
| Fractional width trapezoid | $b$ | Dimensionless | 1 | Arbitrary |
| Proton permeability | $P_{\mathrm{H}^+}$ | m $s^{-1}$ | $10^{-4}$ | [6][1] |
| Hydroxide ion permeability | $P_{\mathrm{OH}^-}$ | m $s^{-1}$ | $10^{-4}$ | Arbitrary |
| Sodium permeability | $P_{\mathrm{C}^+}$ | m $s^{-1}$ | $5\times10^{-10}$ | [29] |
| Chloride permeability | $P_{\mathrm{A}^-}$ | m $s^{-1}$ | $2\times10^{-10}$ | [29] |

**Table S2.** Constant parameters.

We obtained $\Delta G_{\mathrm{NADH}}$ from eQuilibrator [11], which gives the Gibbs free energy of the half-reaction $\mathrm{NADH(aq)} \rightleftharpoons \mathrm{NAD^+(aq)} + 2e^-$ as -64.1 kJ/mol and of the half-reaction $O_2\mathrm{(aq)} + 4e^- \rightleftharpoons 2H_2O$ as $-314.5$ kJ/mol at neutral pH and 100 mM ionic strength. Given that

$$
\frac{1}{2}O_2 + \mathrm{NADH} + \mathrm{H}^+ \rightleftharpoons H_2O + \mathrm{NAD}^+ \tag{S51}
$$

then $\Delta G_{\mathrm{NADH}}{=}-64.1 - 314.5/2 = 221.3$ kJ/mole, which we convert to volts. Similarly, we use the standard Gibbs free energy for ATP hydrolysis of $\Delta G^\circ_{\mathrm{ATP}} = -37.6$ kJ/mol [30] and assume that in *E. coli* $[\mathrm{ATP}] = 10$ mM, $[\mathrm{ADP}] = 0.6$ mM, and $[\mathrm{Pi}] = 20$ mM [9]. Note that these values were reported for *E. coli* growing aerobically on glucose and are slightly higher, but of the same order of magnitude, as the values (2 mM) obtained from single cell experiments also on *E. coli* grown aerobically on glucose [31].

Thus, $\Delta G_{\mathrm{ATP}} = \Delta G^\circ_{\mathrm{ATP}} + RT\ln([\mathrm{ATP}]/[\mathrm{ADP}]/[\mathrm{Pi}]) = -37600 + 8.31\times298\times\ln(0.01/0.02/0.0006) \approx -54$ kJ/mol, which we convert to volts (Table S2). From these values of $\Delta G_{\mathrm{NADH}}$ and $\Delta G_{\mathrm{ATP}}$, we calculate the physiological values of the PMF for the respirative and fermentative regimes (Table S1).

***Minimising the cost.*** Specifying the osmotic pressure, the PMF, intra- and extracellular pH, and the concentrations of all other extracellular ions, we define the cost of maintaining this steady state as the flux of protons that are neither directly involved in generating ATP during respiratory growth nor are actively pumped by $\mathrm{F_1F}_o$ during fermentative growth:

$$\mathrm{Cost} = j_{\mathrm{H}^+} - j_{\mathrm{OH}^-} + 3j_{\mathrm{NhaB}} + 2j_{\mathrm{NhaA}} + j_{\mathrm{ClcA}} \tag{S52}$$

First, we emphasize that now all antiporters with equivalents in *E. coli*, NhaA, NhaB, ClcA, can pump simultaneously, where anions are pumped only by ClcA:

$$\frac{d[A^-]_i}{dt} = j_{\mathrm{A}^-} - 2j_{\mathrm{ClcA}} \tag{S53}$$

The steady-state equation for $\mathrm{A}^-$ is

$$j_{\mathrm{A}^-} = 2j_{\mathrm{ClcA}} \tag{S54}$$

or

$$j^+_{\mathrm{A}^+}\left(1 - e^{\dfrac{F}{RT}\Delta G_{\mathrm{A}^-}}\right) = 2j^+_{\mathrm{ClcA}}\left(1 - e^{\dfrac{F}{RT}(\Delta G_{\mathrm{H}^+} - 2\Delta G_{\mathrm{A}^-})}\right) \tag{S55}$$

We know from Eq.10 that $\Delta G_{H^+} < 0$, thus Eq. S54 requires both sides to be positive for a solution to exists, and hence

$$\frac{1}{2}\Delta G_{\mathrm{H}^+} \leq \Delta G_{\mathrm{A}^-} \leq 0 \tag{S56}$$

Similarly, we require

$$2\Delta G_{\mathrm{H}^+} \leq \Delta G_{\mathrm{C}^+} \leq 0 \tag{S57}$$

for there to be a steady-state solution for cations, because the steady-state equation for $\mathrm{C}^+$ is

$$j_{\mathrm{C}^+} = j_{\mathrm{NhaA}} + 2j_{\mathrm{NhaB}} \tag{S58}$$

or

$$j^+_{\mathrm{C}^+}\left(1 - e^{\dfrac{F}{RT}\Delta G_{\mathrm{C}^+}}\right) = j^+_{\mathrm{NhaA}}\left(1 - e^{\dfrac{F}{RT}(2\Delta G_{\mathrm{H}^+} - \Delta G_{\mathrm{C}^+})}\right) + 2j^+_{\mathrm{NhaB}}\left(1 - e^{\dfrac{F}{RT}(3\Delta G_{\mathrm{H}^+} - 2\Delta G_{\mathrm{C}^+})}\right) \tag{S59}$$

Again, we note that Eq. S58 requires Eq. S57 for a solution.

Second, from Eq. S13, $[\mathrm{H}^+]_i$, and $[\mathrm{H_2O}]_i = 55.56$ M, we calculate $[\mathrm{OH}^-]_i$ at steady state, assuming that the leakage of hydroxide ions is described by Eq. S9. Then both $j_{\mathrm{H}^+}$ and $j_{\mathrm{OH}^-}$ in Eq. S52 are known. Specifically, we use here Erying's equation Eq. S10 to compute $j_{\mathrm{H}^+}$ and $j_{\mathrm{OH}^-}$ but could have used for instance GHK equation. Further, Eqs. S54 and S58 allow us to rewrite the cost in two ways:

$$\mathrm{Cost} = j_{\mathrm{H}^+} - j_{\mathrm{OH}^-} + \frac{1}{2}j_{\mathrm{A}^-} + \begin{cases} 2j_{\mathrm{C}^+} - j_{\mathrm{NhaB}} \\ \frac{3}{2}j_{\mathrm{C}^+} + \frac{1}{2}j_{\mathrm{NhaA}} \end{cases} \tag{S60}$$

either in terms of $j_{\mathrm{NhaB}}$ or $j_{\mathrm{NhaA}}$.

Third, we combine the equation for the osmotic pressure

$$\frac{\Delta\Pi}{RT} = [Y]_i + [\mathrm{H}^+]_i - [\mathrm{H}^+]_e + [\mathrm{OH}^-]_i - [\mathrm{OH}^-]_e - [\mathrm{C}^+]_e\left(1 - e^{\dfrac{F}{RT}(\Delta G_{\mathrm{C}^+} - \Delta\psi)}\right) - [\mathrm{A}^-]_e\left(1 - e^{\dfrac{F}{RT}(\Delta G_{\mathrm{A}^-} + \Delta\psi)}\right) \tag{S61}$$

with the equation for the membrane potential

$$\frac{SC_m}{FV}\Delta\psi = z_Y[Y]_i + [\mathrm{H}^+]_i - [\mathrm{OH}^-]_i + [\mathrm{C}^+]_e \cdot e^{\dfrac{F}{RT}(\Delta G_{\mathrm{C}^+} - \Delta\psi)} - [\mathrm{A}^-]_e \cdot e^{\dfrac{F}{RT}(\Delta G_{\mathrm{A}^-} + \Delta\psi)} \tag{S62}$$

to express $\Delta G_{\mathrm{C}^+}$ in terms of $\Delta G_{\mathrm{A}^-}$ and $z_Y$, with all the other variables being specified.

We are now able to minimise the cost for these conditions. For negative membrane potential, and given Eq. S56, we simultaneously scan $\Delta G_{\mathrm{A}^-}$ over this range and $z_Y$ from $-10 \leq z_Y \leq 10$. For each value of $\Delta G_{\mathrm{A}^-}$ and $z_Y$, we find $\Delta G_{\mathrm{C}^+}$ from Eqs. S61 and S62. For these choices to be steady state, then additionally we require both that $[Y]_i$, found from Eq. S61, is not negative and that Eq. S57 holds.

For those values of $\Delta G_{\mathrm{A}^-}$, $z_Y$, and $\Delta G_{\mathrm{C}^+}$ that satisfy these requirements and do allow a steady state, then $j_{\mathrm{A}^-}$ and $j_{\mathrm{C}^+}$ in Eq. S60 are known and computed according to Erying's equation Eq. S10. We need consider only two further cases:

- If the calculated $\Delta G_{\mathrm{C}^+}$ is such that $\frac{3}{2}\Delta G_{\mathrm{H}^+} \leq \Delta G_{\mathrm{C}^+} \leq 0$, then both $j_{\mathrm{NhaA}}$ and $j_{\mathrm{NhaB}}$ are positive from Eq. S59. The cost is minimised if $j^+_{\mathrm{NhaA}}$ is zero, using the lower branch of Eq. S60, and this minimal cost is given by Eq. S60 with $j_{\mathrm{NhaA}} = 0$.
- If the calculated $\Delta G_{\mathrm{C}^+}$ is such that $2\Delta G_{\mathrm{H}^+} \leq \Delta G_{\mathrm{C}^+} \leq \frac{3}{2}\Delta G_{\mathrm{H}^+}$, then $j_{\mathrm{NhaA}}$ is again positive but $j_{\mathrm{NhaB}}$ is negative from Eq. S59. The cost is minimised if $j^+_{\mathrm{NhaB}}$ is zero, using the upper branch of Eq. S60, and this minimal cost is given by Eq. S60 with $j_{\mathrm{NhaB}} = 0$.

For each scanned pair of $\Delta G_{\mathrm{A}^-}$ and $z_Y$, we can therefore calculate the cost, and by comparing all pairs, we choose the $\Delta G_{\mathrm{A}^-}$ and $z_Y$ that give the minimal cost. For positive membrane potential we scan over $\Delta G_{\mathrm{C}^+}$ and $z_Y$, in the exact same way as described above for $\Delta G_{\mathrm{A}^-}$ and $z_Y$. In our scans, we explicitly include the case where no antiporters are needed: $\Delta G_{\mathrm{A}^-} = \Delta G_{\mathrm{C}^+} = 0$, which always minimises the cost if it allows a steady state. We find that when antiporters are used, the optimal $z_Y$ is always at an extremum, either $z_Y = 10$ (when $\Delta\psi > 0$) or $z_Y = -10$ (when $\Delta\psi < 0$).

**Approximating the optimal strategy** Interestingly, we find numerically that for an arbitrary range of $z_Y \in [\min(z_Y), \max(z_Y)]$ and an osmotic pressure of 1 atm, the following strategy approximates well the optimal solution (Fig. S15:

1. If the desired membrane potential can be achieved by setting $\Delta G_{\mathrm{C}^+} = \Delta G_{\mathrm{A}^-} = 0$, the strategy is given by:

$$\begin{cases} z_Y & \text{given by Eq. S62} \\ j^+_{\mathrm{ClcA}} & = 0 \\ j^+_{\mathrm{NhaB}} & = 0 \\ j^+_{\mathrm{NhaA}} & = 0 \\ \text{Cost} & = j_{\mathrm{H}^+} - j_{\mathrm{OH}^-} \end{cases} \tag{S63}$$

2. If the desired membrane potential is negative and cannot be achieved by setting $\Delta G_{\mathrm{C}^+} = \Delta G_{\mathrm{A}^-} = 0$, then set:

$$\begin{cases} z_Y & = \min(z_Y) \\ \Delta G_{\mathrm{A}^-} & = 0 \ \text{ and } \ j^+_{\mathrm{ClcA}} = 0 \end{cases} \tag{S64}$$

and compute the corresponding electrochemical potential of cations required to achieve the desired $\Delta\psi$ given zero AMF. Depending on the value of this CMF, the solution is:

$$\text{if } \Delta G_{\mathrm{C}^+} \in [3/2\Delta G_{\mathrm{H}^+}, 0] \quad \text{then} \quad \begin{cases} j^+_{\mathrm{NhaA}} & = 0 \\ j_{\mathrm{NhaB}} & = 2j_{\mathrm{C}^+} \\ \text{Cost} & = j_{\mathrm{H}^+} - j_{\mathrm{OH}^-} + \dfrac{3}{2} j_{\mathrm{C}^+} \end{cases} \tag{S65}$$

$$\text{if } \Delta G_{\mathrm{C}^+} \in [2\Delta G_{\mathrm{H}^+}, 3/2\Delta G_{\mathrm{H}^+}] \quad \text{then} \quad \begin{cases} j^+_{\mathrm{NhaB}} & = 0 \\ j_{\mathrm{NhaA}} & = j_{\mathrm{C}^+} \\ \text{Cost} & = j_{\mathrm{H}^+} - j_{\mathrm{OH}^-} + 2j_{\mathrm{C}^+} \end{cases} \tag{S66}$$

3. If the desired membrane potential is positive and cannot be achieved by by setting $\Delta G_{\mathrm{C}^+} = \Delta G_{\mathrm{A}^-} = 0$, then set:

$$\begin{cases} z_Y & = \max(z_Y) \\ j^+_{\mathrm{NhaB}} = j^+_{\mathrm{NhaA}} & = 0 \ \text{ and } \ \Delta G_{\mathrm{C}^+} = 0 \\ j_{\mathrm{ClcA}} & = 1/2 j_{\mathrm{A}^-} \\ \text{Cost} & = j_{\mathrm{H}^+} - j_{\mathrm{OH}^-} + 1/2 j_{\mathrm{A}^-} \end{cases} \tag{S67}$$

To compute leakage fluxes: $j_{\mathrm{H}^+}$, $j_{\mathrm{OH}^-}$, $j_{\mathrm{C}^+}$ and $j_{\mathrm{A}^-}$, we use Erying's equation Eq. S10. Occasionally the membrane potential required to achieve a given $\mathrm{pH}_i$, PMF, and osmotic pressure is simply not achievable for any state of the system. We then require the membrane potential to be the one with the minimal deviation of the resulting $\mathrm{pH}_i$ from the desired $\mathrm{pH}_i$.

### Supplementary Figures

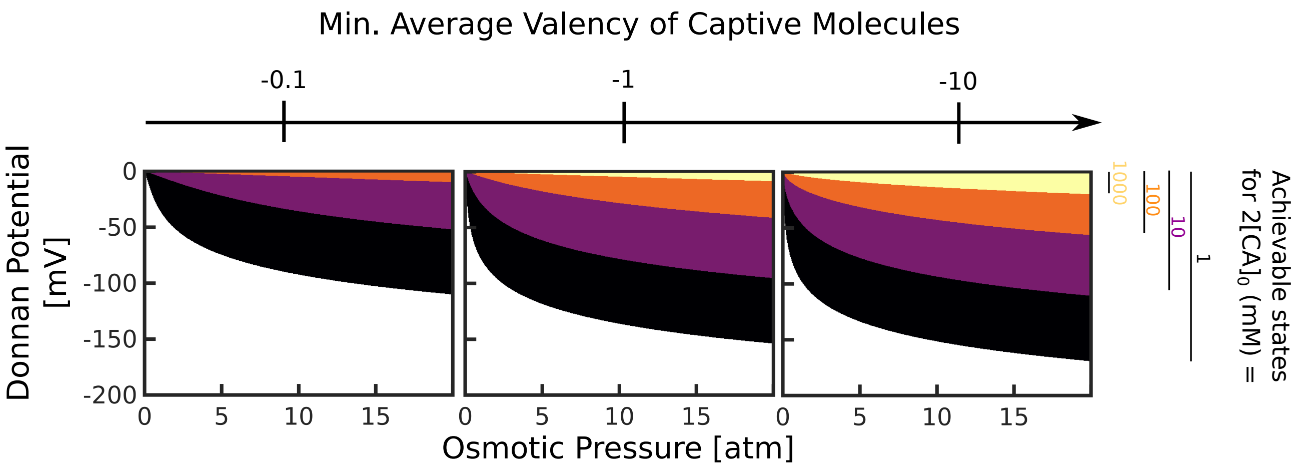


**Fig. S1. Membrane potential can be maintained in thermodynamic equilibrium – Donnan potential – by captive charged molecules alone, but require low salinities of the medium, large average valencies of captive charges, and elevated osmotic pressures.** We set pH$_i$ $=$ pH$_e$ $= 7$, $\Delta G_{C+} = \Delta G_{A-} = 0$ and show the steady-state $\Delta\psi$ and $\Delta\Pi$ given a salinity [CA]$_0$ $\in \{0.5, 5, 50, 500\}$ mM and a minimal average valency of captive molecules. We note that the steady states achievable by [CA]$_0$ $= 0.5$ mM also includes those achievable for any value of [CA]$_0$ $> 0.5$ mM – all four coloured regions are achievable by setting [CA]$_0$ $= 0.5$ mM.

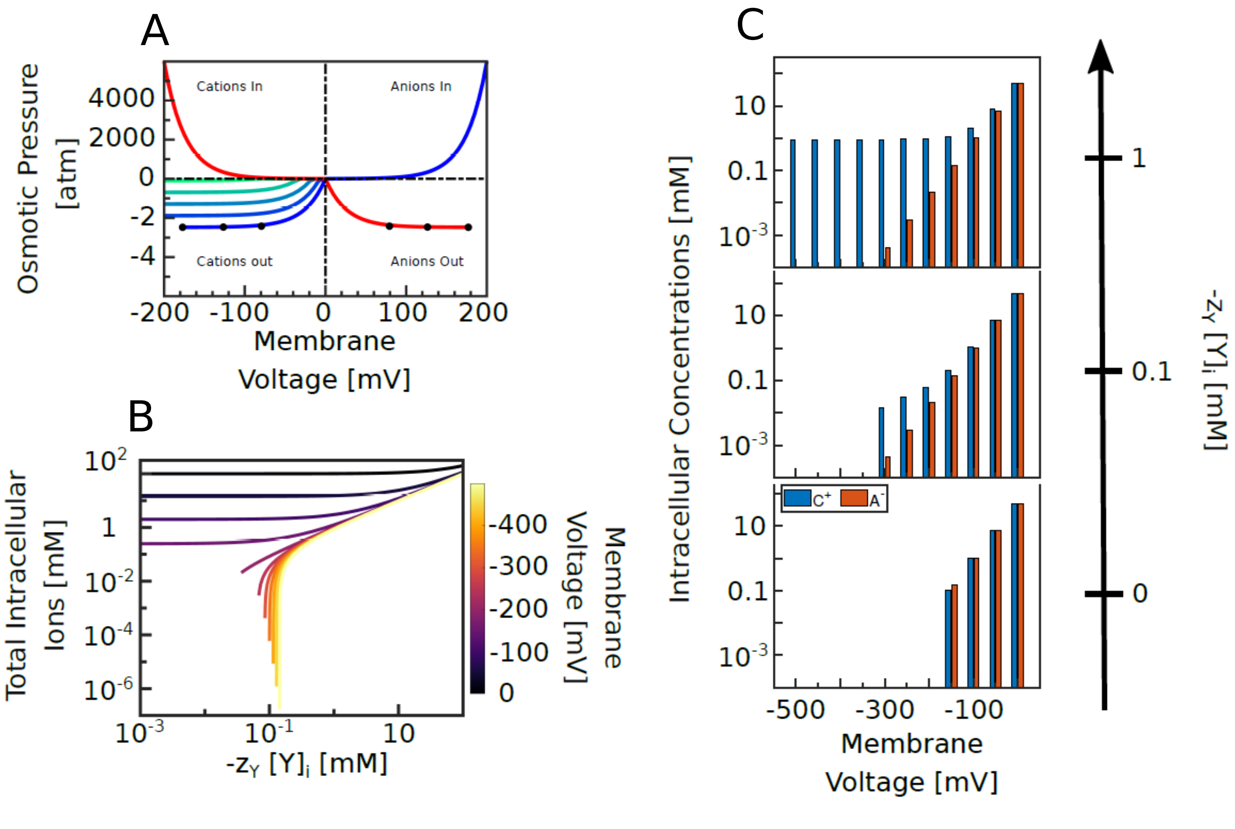


**Fig. S2. Strategies for generating physiological $\Delta\psi$ and $\Delta\Pi$** (A) Osmotic pressure values (calculated only based on ion concentrations) plotted against $\Delta\psi$ when one cation (blue), or one anion (red), are pumped either in or out of the cell. To obtain the plotted $\Delta\psi$ values we set pH$_i$ = 7. When only anions are pumped we set $\Delta G_{C+} = 0$ and vary $\Delta G_{A-}$ between $\pm$500 mV, and similarly, when cations only are pumped we set $\Delta G_{A-} = 0$ and vary $\Delta G_{C+}$ between $\pm$500 mV. [Y]$_i$ = 0 in all but the bottom left, where darkest blue indicates [Y]$_i$ =0 and lines from dark blue to light blue [Y]$_i$= 0, 12, 24, 36 and 48 % of the total extracellular ionic concentration ([CA]$_0$). Black dots on the dark blue line in the bottom left, and the red line in the bottom right indicate the minimum/maximum $\Delta\psi$ that can be reached for total extracellular ion concentrations of 1, 10 and 100 mM (the dot for 1 mM is the one closest to $\Delta\psi$ = 0). For all the other lines in the figure, [CA]$_0$=100 mM. Apart from the minimum/maximum value of $\Delta\psi$, changing the total extracellular ion concentration will not change $\Delta\psi$ behaviour. (B) $\Delta\psi$ (for cations pumped out) is composed of the contribution from the captive charges (x axis, only negative $z_Y[Y]_i$ are considered and the absolute value is plotted) and intracellular ions contribution (y axis). We set pH$_e$ = pH$_i$ = 7; [CA]$_0$ = 100 mM. Cells that harbour small amounts of $z_Y[Y]_i$ become increasingly depleted of small ions as they become more depolarised (the absolute value of $\Delta\psi$ increases). Since $\Delta\psi$ is maintained by pumping cations out, it becomes impossible to depolarize the cell further when $[C]_i \to 0$. (C) Intracellular ionic composition at different $\Delta\psi$ and for three values of $z_Y[Y]_i$. Pumping cations out results in [A]$_i$ > [C]$_i$ when $z_Y[Y]_i$ = 0 and [C]$_i$ > [A]$_i$ when $z_Y[Y]_i$ < -0.1 mM.

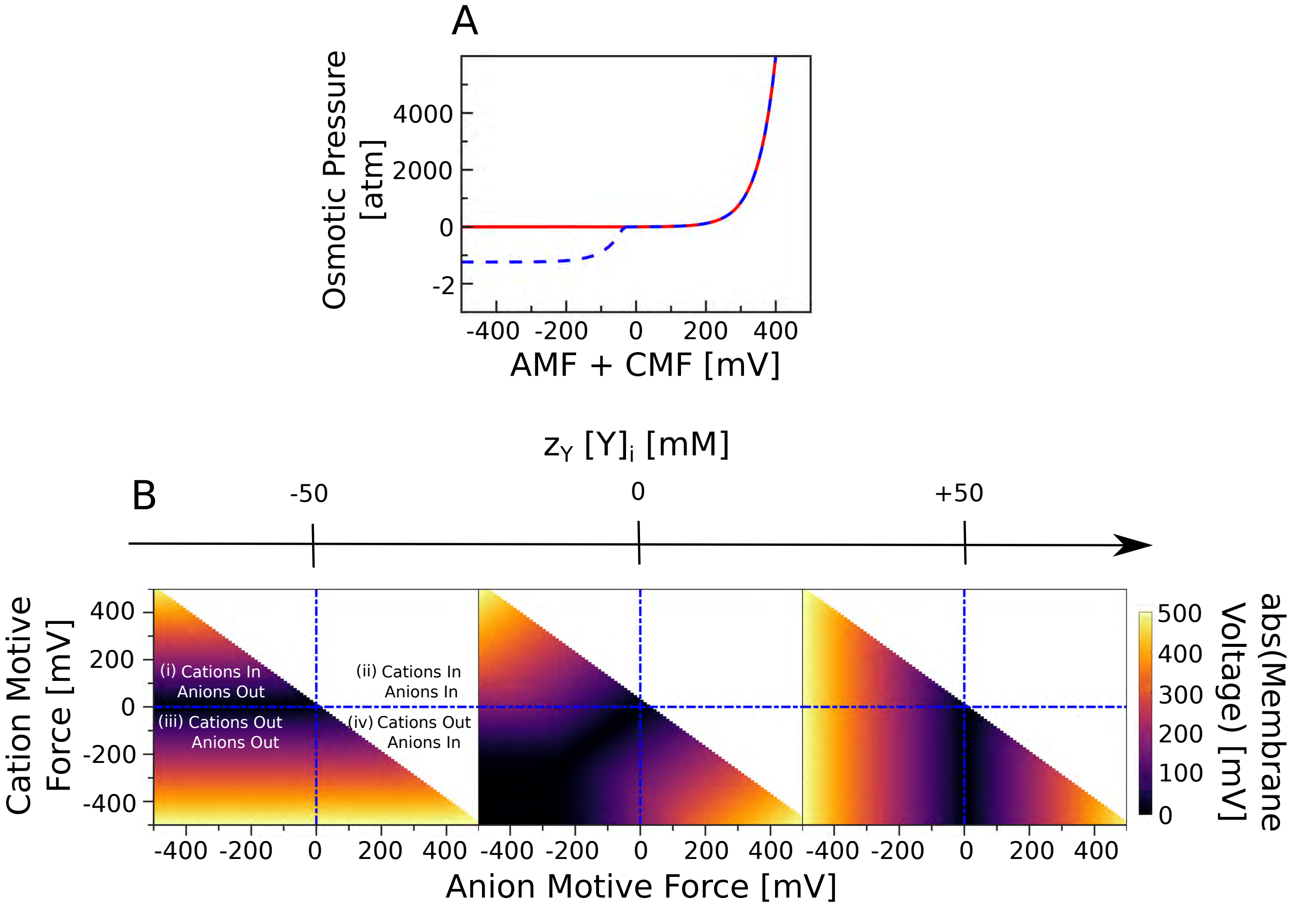


**Fig. S3. Membrane voltage and osmotic pressure generated when both small cations and anions are pumped.** We keep [CA]$_0$ = 50 mM, pH$_i$ = pH$_e$ = 7. (A) Relationship between osmotic pressure and AMF + CMF. Red and blue are for $z_Y[Y]_i$ respectively 0 and ±50 mM. Keeping the osmotic pressure in check translates to $\Delta G_{C+} + \Delta G_{A-} \leq 0$. The osmotic pressure is only a function of $z_Y[Y]_i$ and the sum of the ionic motive forces, irrespective of their relative values. (B) Absolute value of $\Delta\psi$ (colour map) is plotted for different CMF and AMF values; it is positive north-west and negative south-east. Only simulations resulting in $\Pi < 3$ atm are shown. $\Delta\psi$ is not sensitive to the AMF when $z_Y[Y]_i$ = -50 mM. Similarly, $\Delta\psi$ is insensitive to CMF when $z_Y[Y]_i$ = 50 mM.

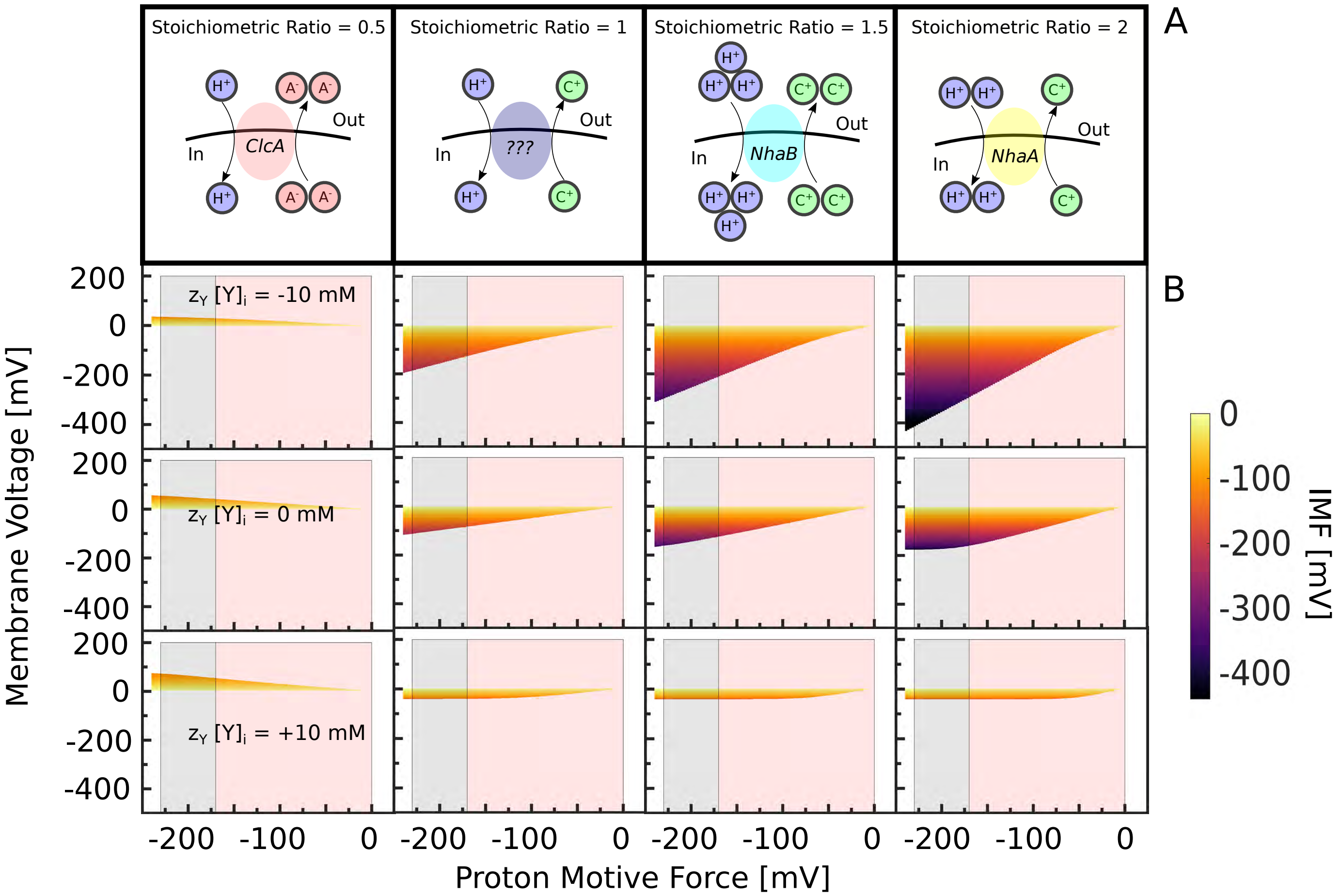


**Fig. S4. The steady-state membrane potential required to maintain a neutral pH$_i$ for a given PMF.** (A) Schemes of the antiporters we study. (B) We set $[\text{CA}]_0 = 100$ and pH$_i = 7$ and show the range of membrane potential ($y$-axis) that can be maintained at steady state for each PMF ($x$-axis), as well as the corresponding IMF (colour scale). Each value of the membrane potential corresponds to a value of the extracellular pH shown in Fig. 1D. The grey area indicates the PMF in the respirative regime; the pink area the fermentative regime.

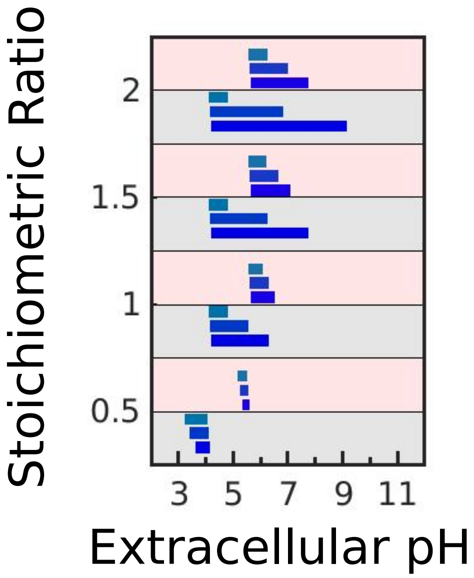


**Fig. S5. Altering $z_Y[Y]_i$ and the PMF changes the range of extracellular pH over which *E. coli* is able to maintain a neutral pH$_i$.** Following the approach of Fig. 1D, we show the predicted pH$_e$ ranges for two values of the PMF: -170 mV for the respirative regime in grey and -85 mV for the fermentative regime in pink. Each of the four antiporters of Fig. 1C are indicated by their stoichiometric ratio. We consider three different values of $z_Y[Y]_i$: -10 mM, 0, and +10 mM, shown using deep to light blue.

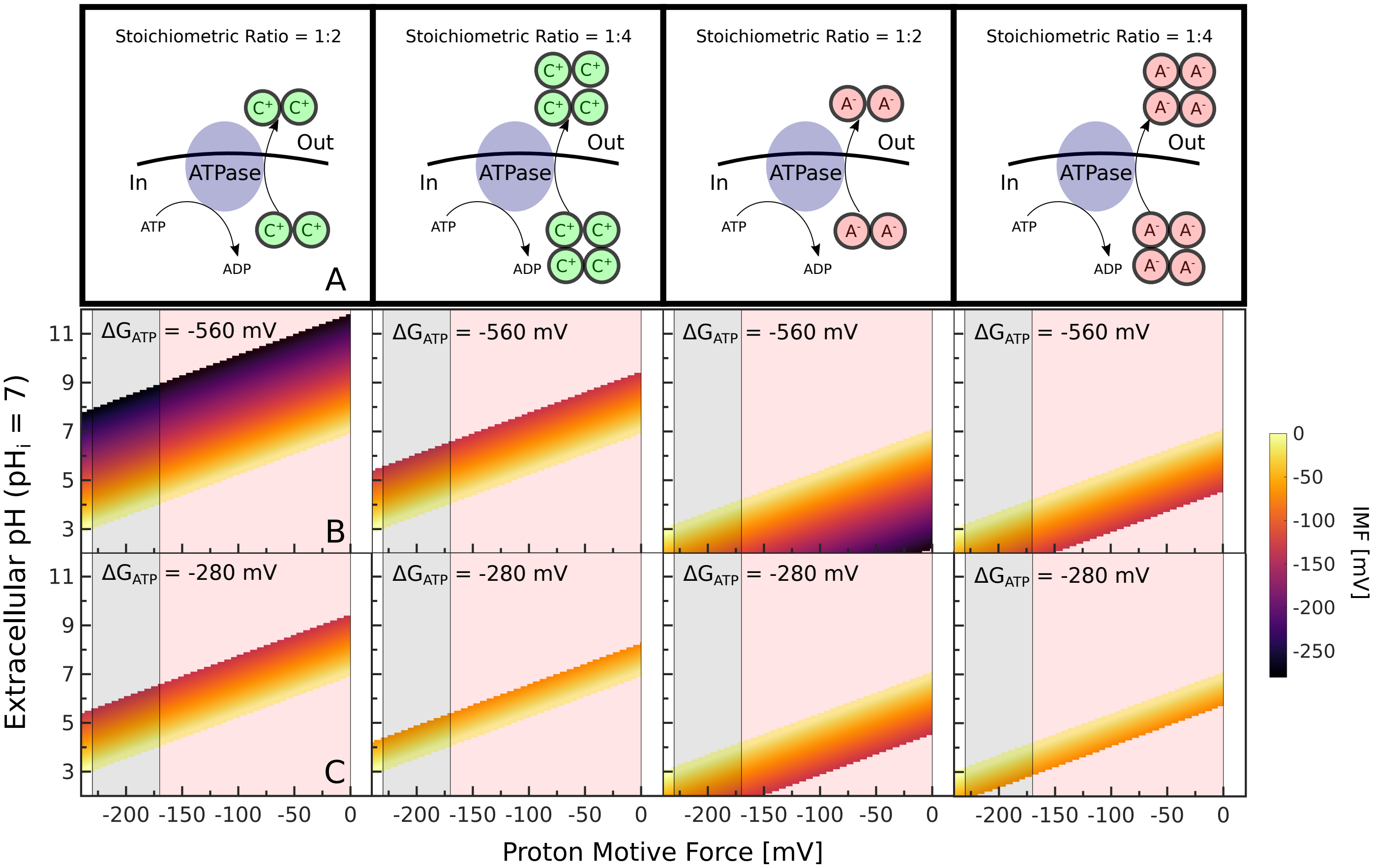


**Fig. S6. The pH$_e$ range over which cells can maintain a neutral pH$_i$ does not scale with PMF if ATP-driven efflux pumps are used for generating physiological membrane potential.** (A) Schematics of hypothetical ATP-driven ion efflux pumps we consider. (B) and (C) We plot the possible steady-state solutions for the hypothetical pumps for a given PMF between -240 and 0 mV, a pH$_i$ of 7, and pH$_e$ between 2 and 12. The average valency of captive molecules is bound between -1 and +1, osmotic pressure is set to 1 atm, and extracellular [CA]$_0$ = 100 mM. Grey and red shading shows values of the PMF expected for respiration and fermentation. The colour scale indicates the ionic motive force (IMF). (B) shows the case where the free energy of ATP hydorlysis is -560 mV and (C) -280 mV. Halving $\Delta G_{ATP}$ is equivalent to halving the stoichiometric ratio of the ATP efflux pump. Contrary to the case when ion efflux is powered by the PMF, i.e. is a consequence of the action of proton:ion antiporters, the range of pH$_e$ at which pH$_i$ can be maintained does not scale with the PMF. Both ways of powering ion efflux however result in loss of pH$_i$ homeostasis at acidic pH$_e$ upon loss of PMF and when the cell pumps anions, and the ability to maintain pHi upon loss of PMF is impaired more when efflux is powered by PMF.

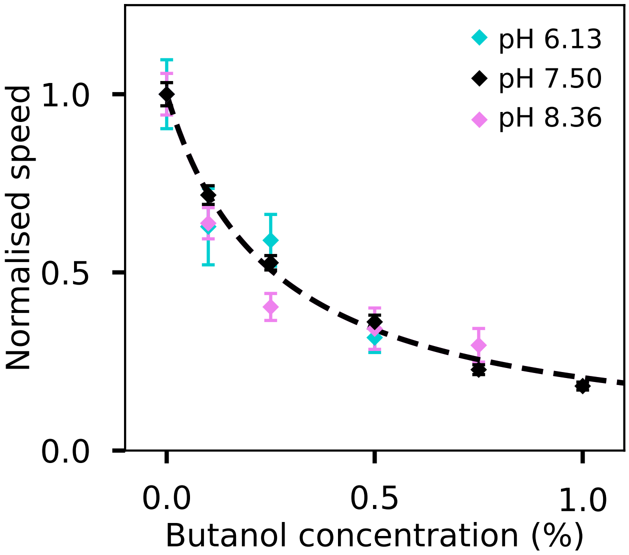


**Fig. S7. The bacterial flagellar motor speed remains proportional to PMF within pH$_i$ 6.1 to 8.4 range.** Bacterial flagellar motor speed has been found proportional to the PMF at different loads [32, 33]. Here we confirm that (at our chosen motor load) the motor speed remains proportional to PMF at different pH$_i$. For this purpose, *E. coli* cells were grown and prepared as described in the main text *Methods* section. To control pH$_i$ cells in our tunnel slide were exposed to the motility buffer (BMB, see *Methods* in the main text) of pH 6.13 or 8.36 containing 40 mM potassium benzoate and 40 mM methylamine hydrochloride, which equilibrates internal and external pH [34]. Next, we exposed the cells with pH$_i$ 6.13 and 8.36 to different butanol concentrations, where we know how the motor speed responds to different butanol concentrations at near neutral pH$_i$ from our previous studies [35]. The results are plotted together; previous results are in black, pink are the results for pH$_i$=6.13 and cyan 8.36. The black line shows a hyperbola fit also taken from [35] and the error bars show the standard error of the mean. Normalised speeds from different cells are plotted against the butanol concentration for each pH$_i$ (total of 24 cells for pH$_i$ 6.13 and 40 for 8.36). Normalised motor speeds follow the same trendline irrespective of pH$_i$.

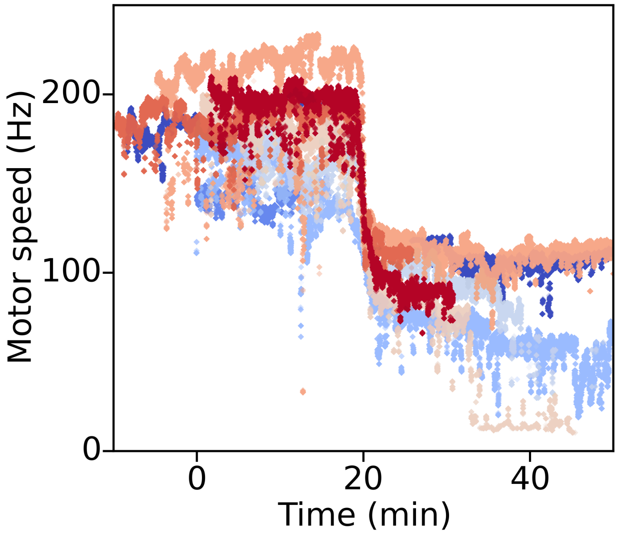


**Fig. S8. In the absence of oxygen, the measured PMF approximately halves.** Single-motor traces of *E. coli* cells (eight plotted using a different colour) in the sealed tunnel-slides at pH$_e$=7.0 in the presence of glucose. Cells are grown and prepared as described in the main text *Methods* section. Briefly upon harvesting cells are washed into BMB pH 7.0, supplemented with 20 mM glucose, and attached to the tunnel slide. Fresh BMB is flushed through the tunnel shortly before the recording, upon which the tunnel is sealed with a CoverGrip™ Coverslip Sealant to prevent oxygen exchange. The recording continues for 30-50 min. The traces are aligned by the time at which the motor speed drops – here shown at 20 min. The actual time point varied between 20 and 30 min depending on the concentration of cells and the period between sealing the tunnel-slide and beginning the recording. Speed drop indicates oxygen depletion in the medium, as reported in [36]

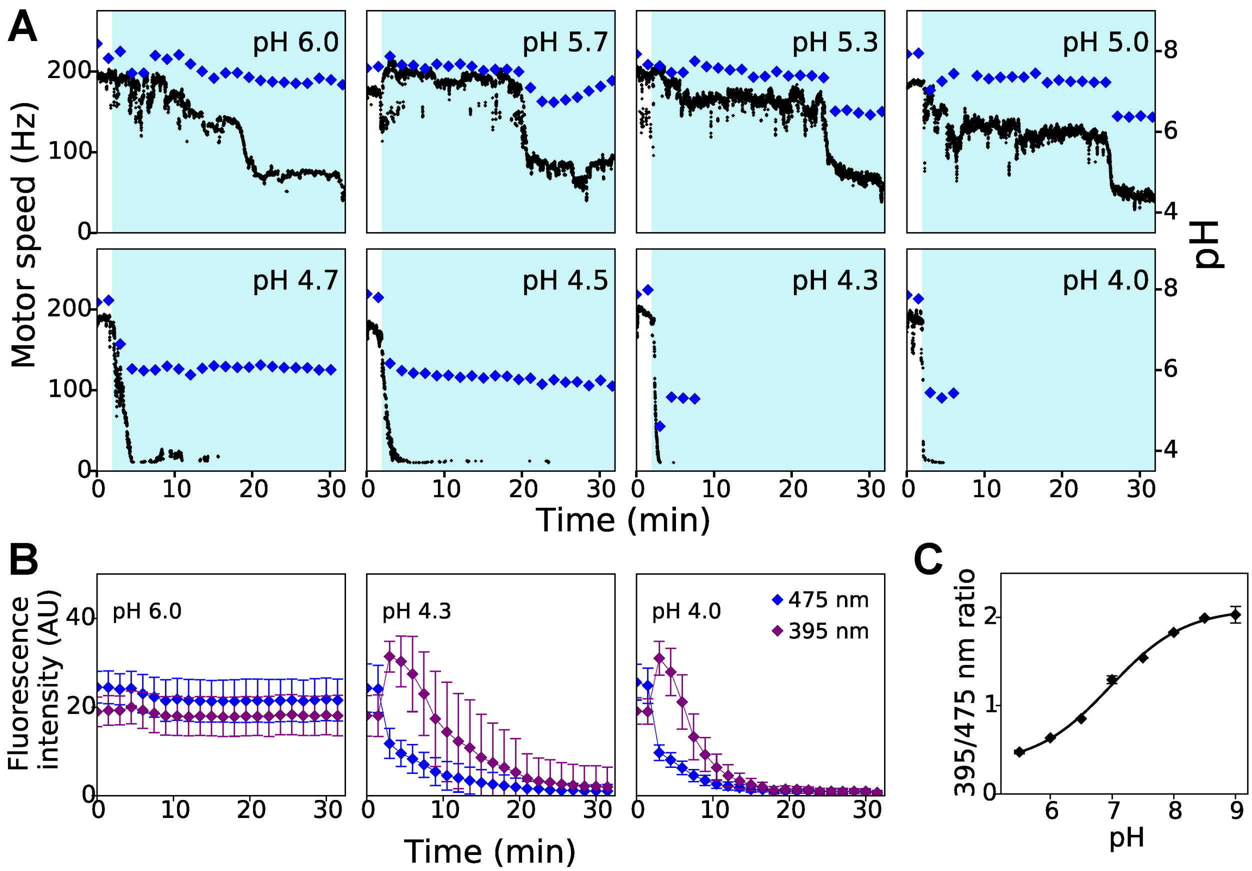


**Fig. S9. Cells maintain neutral pH$_i$ at pH$_e$ $\geq$ 5.0 for high magnitude PMF (i.e. in the presence of oxygen), and only at pH$_e$ $\geq$ 5.5 for lower PMF (in anaerobic conditions).** (A) Single-cell examples of speed (in black) and pH$_i$ (in blue) dynamics at different pH$_e$. Cells are prepared as described in the main text *Methods* section. At time t=0 to 2 min cells are in BMB, supplemented with 20 mM glucose at pH$_e$=7 (white background); 2 min after the recording commences, low pH buffers are introduced (cyan region) and the slide is sealed. Sharp motor speed drop observed in each example trace indicates the transition from aerobic to anaerobic metabolism, which happens 20-30 minutes after sealing the slide as shown also in S8 and in [36]. As the speed drops, pH$_i$ drops as well. For pH$_e$ $\geq$ 5.5, pH$_i$ recovers to neutral even after the PMF loss, however, for 5.0$\leq$ pH$_e$ $\leq$ 5.5 pH$_i$ is maintained neutral only at higher motor speeds (i.e. in aerobic conditions) but not after oxygen is depleted (B) pHluorin is not a suitable sensor for low pH$_i$ because fluorescence is lost at low pH$_e$ at both the relevant wavelengths of 475 nm and 395 nm. For each wavelength, we plot the averaged relative fluorescence intensity of $\approx$ 30 cells, grown and prepared for imaging as described in the main text *Methods* section that also contains the imaging conditions. Error bars are the standard deviation of the pixels' intensities. (C) The sensitivity range of pHluorin spans pH 6–8.5. The calibration curve is reproduced from [37], where the pH gradient between the medium and the cytoplasm is collapsed by addition of 40 mM potassium benzoate and 40 mM methylamine hydrochloride and the buffers of known pH are used to calibrate pHluorin spectrum changes (see also *Methods* section in the main text). The mean 395/475 intensity ratio is plotted with the standard deviation against pH and fitted with a sigmoid function. Any pH$_i$ lower than 5.5 will be detected as pH$_i$=5.5.

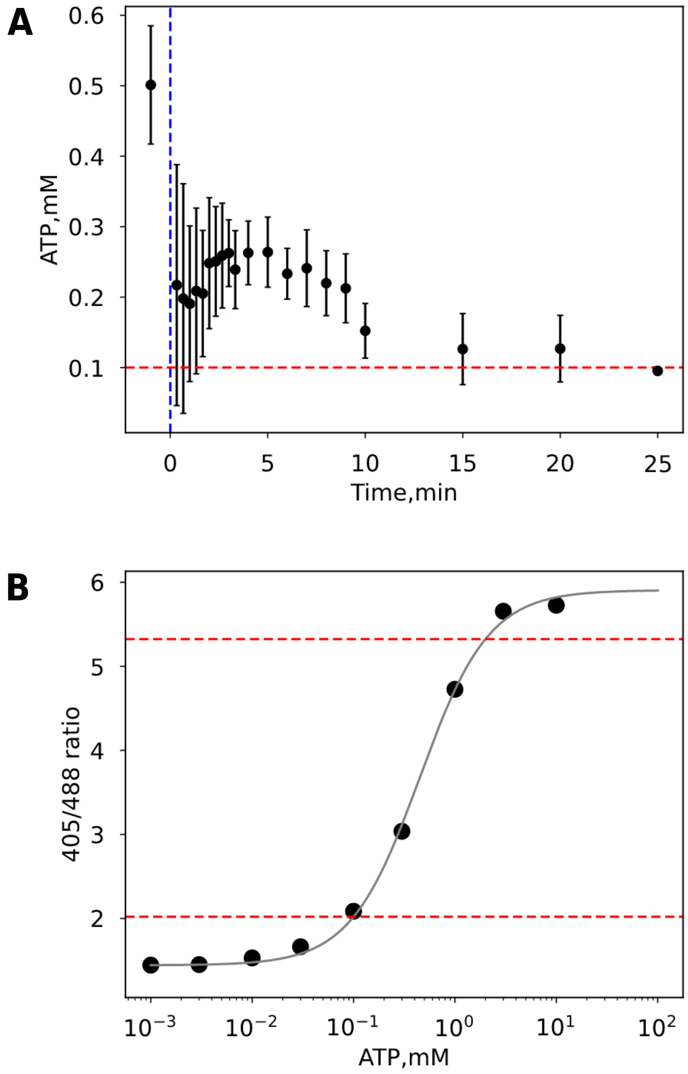


**Fig. S10. *E. coli* maintain non-zero ATP levels for ∼10 min after the addition of 100 μM CCCP.** (A) ATP concentration in *E. coli* cells measured with a fluorescent ratiometric Queen7μM* sensor [38], a modified version of Queen7μM [31]. Cells, carrying pWR-Q7* plasmid [38] are grown and prepared in BMB buffer with 20 mM glucose and pH 8.13 to 8.2 as described in the main text *Methods.* Fluorescence intensity is measured in the plate reader as described in the main text *Methods* with excitation at 405 and 488 nm and emission at 528 nm, and coverted to [ATP] using the callibration curve in (B). The first point is taken before CCCP addition. At time 0 (blue dashed line), 100 μM CCCP is added to the cells. Red dashed line shows the sensitivity limit of the Queen7μM* sensor, and error bars show the standard deviation of the mean between 4 replicates. (B) Calibration curve for Queen7μM* is obtained as in [38] using lysates of Queen7μM*-expressing cells and known ATP concentrations (see main text *Methods* section). Sensitivity range of the sensor spans from ∼0.1 to ∼2 mM ATP (red dashed lines).

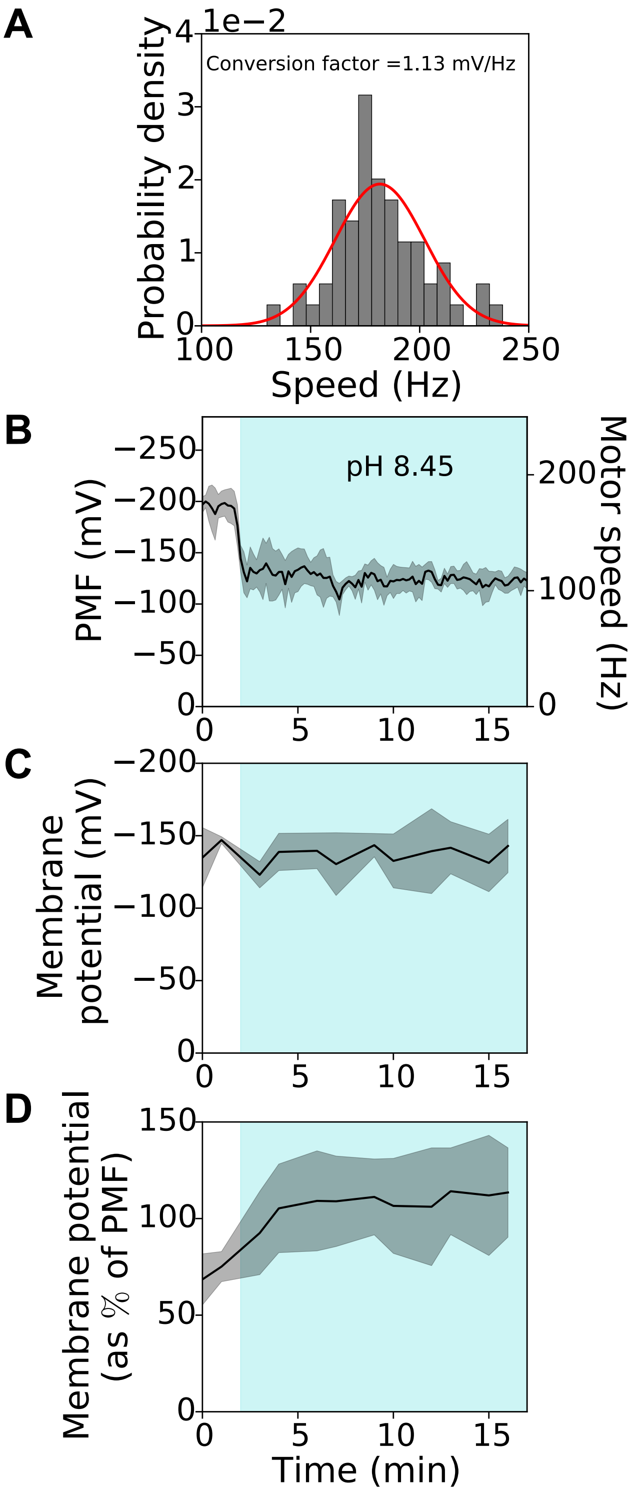


**Fig. S11. Calibration of the PMF measured from the bacterial flagellar motor speeds.** (A) The motor speed in BMB (see *Methods* in the main text) at pH$_e$=7.0 and in the presence of 20 mM glucose is converted to PMF using the predicted PMF range for cells respiring on glucose: $170 \leq$ |PMF| $\leq 229$ mV. We combine 60 single-cell (single-motor) traces collected at pH$_e$ = 7 and computed from 90 s long recordings. Cells were grown and prepared for tunnel slide recording as described in *Methods* section of the main text. The histogram of the resulting motor speeds is plotted – its median is 180.6 Hz; the red line shows a fit to a Gaussian distribution. We calculate the conversion factor that maximises the amount of traces within the predicted PMF range as $(170+229)/2/180.6 \approx 1.1$ mV/Hz. (B) The mean PMF of four cells, grown and prepared for imaging as described in the main text *Methods* section, experiencing an alkaline shift. Cell are kept in BMB (*Methods* in main text) with 20 mM glucose and pH$_e$=7 before the shift (white region), and pH$_e$=8.45 after the shift (cyan region). The PMF is calculated from the time series of motor speeds using the conversion factor from A. (C) The membrane potential and (D) the membrane potential relative to the PMF for the same data. The membrane potential is calculated from Eq. 2, where $\Delta$G$_{H+}$ is the PMF taken from B and the proton concentrations are found by measuring pH$_i$ and pH$_e$. Errors are standard deviations.

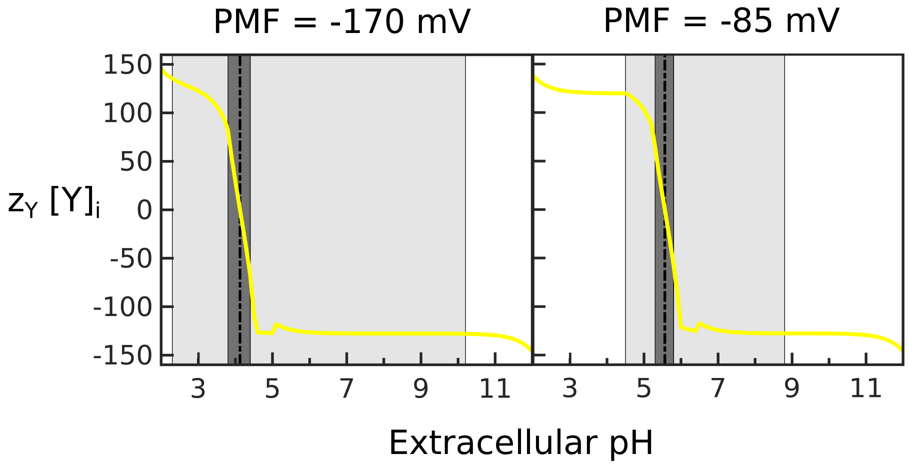


**Fig. S12. The optimal valency changes sign as the pH$_e$ changes** The numerical experiment is identical to that of Fig. 4 in the main text. We maintain pH$_i$ as closed as possible to $7$, $-10 \leq z_Y \leq +10$, set the osmotic pressure to 1 atm and $[\text{CA}]_0$ to 50 mM. The black dotted line marks the pH$_e$ for which $\Delta\psi = 0$; to the left $\Delta\psi > 0$; to the right $\Delta\psi < 0$. The dark grey area indicates where the optimal solution has $\Delta G_{C+} = \Delta G_{A-} = 0$. The light grey area indicates where pH$_i$ can be maintained exactly at 7. For two different PMF values we show $z_Y[Y]_i$ that minimizes the cost of maintaining a steady-state (Eq. 18 in the main text).

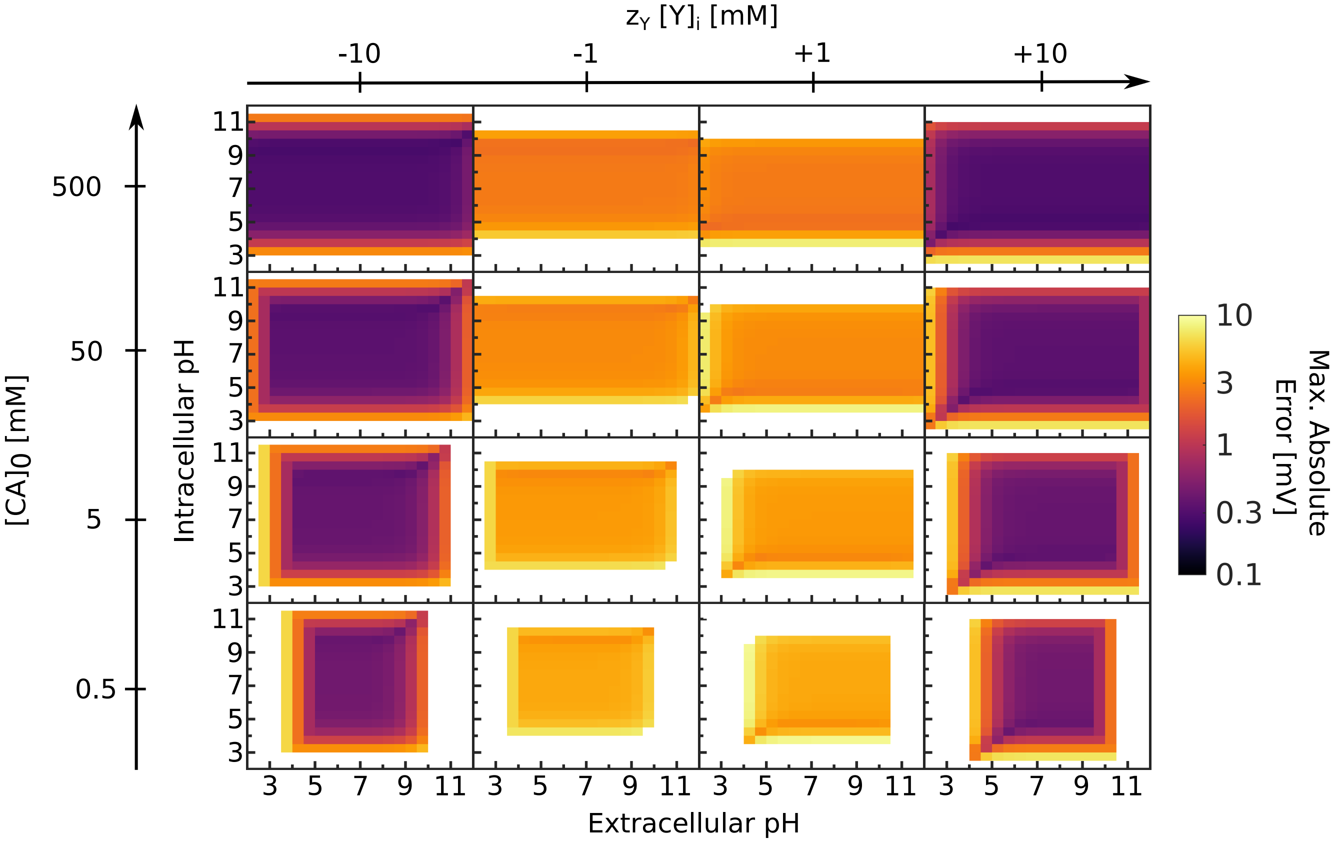


**Fig. S13. Approximating the membrane potential gives low relative errors.** We obtained $\Delta\psi$ in Fig. 1D by solving Eq. S41 for a fixed value of [CA]$_0$=50 mM. Here we plot the maximal absolute error in membrane potential if we assume [CA]$_0$ is large and pH$_e$=7 and calculate $\Delta\psi$ using Eq. Eq. S46 instead. For a given $z_Y[Y]_i$, [CA]$_0$, pH$_e$ and pH$_i$, we find the error for $\Delta G_{C_+}$ and $\Delta G_{A_-}$ varying over an $11 \times 11$ grid of equally spaced values within [-500,+500] mV, but show the maximal absolute error. White regions correspond to maximal errors greater than 10 mV. For $z_Y[Y]_i = 0$ (not shown), the approximation always gives an absolute error greater than 10 mV.

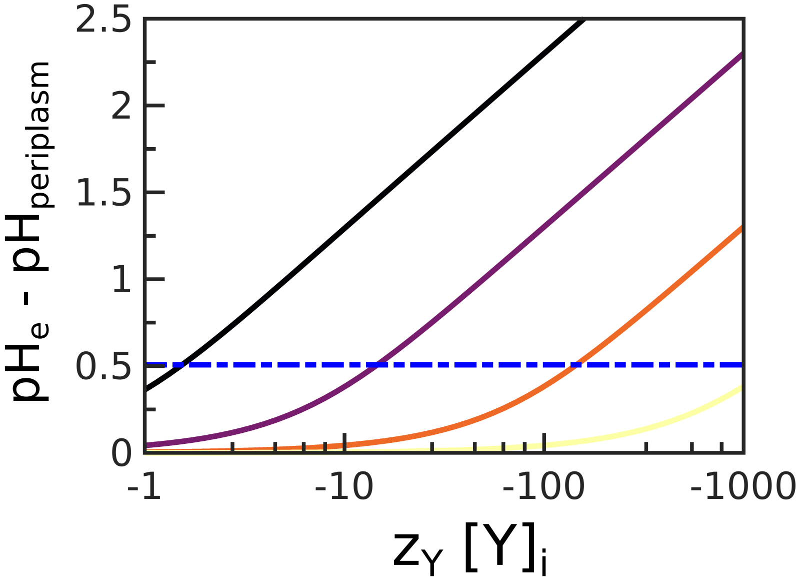


**Fig. S14. The periplasmic membrane potential can offset the pH$_e$ 'felt' by the inner membrane.** The difference between pH$_e$ and the periplasmic pH is plotted as a function of the periplasmic captive charges, $z_Y[Y]_p$, and for different values of [CA]$_0$: black to yellow show 1, 10, 100, and 1000 mM. Using the measured periplasmic radius, *E. coli*'s' periplasmic volume was estimated to be between 8 and 16$\%$ of the total cell volume [39] – we assume a value of $10\%$, and here do not consider osmotic pressure. In addition, we set pH$_e$=7 and let the periplasm be in ionic equilibrium with the environment. At low salinity (black and purple lines), the periplasmic captive charges offset the pH$_e$ 'observed' by the inner membrane. The difference between pH$_e$ at the inner membrane decreases with [CA]$_0$ even with little periplasmic captive charges.

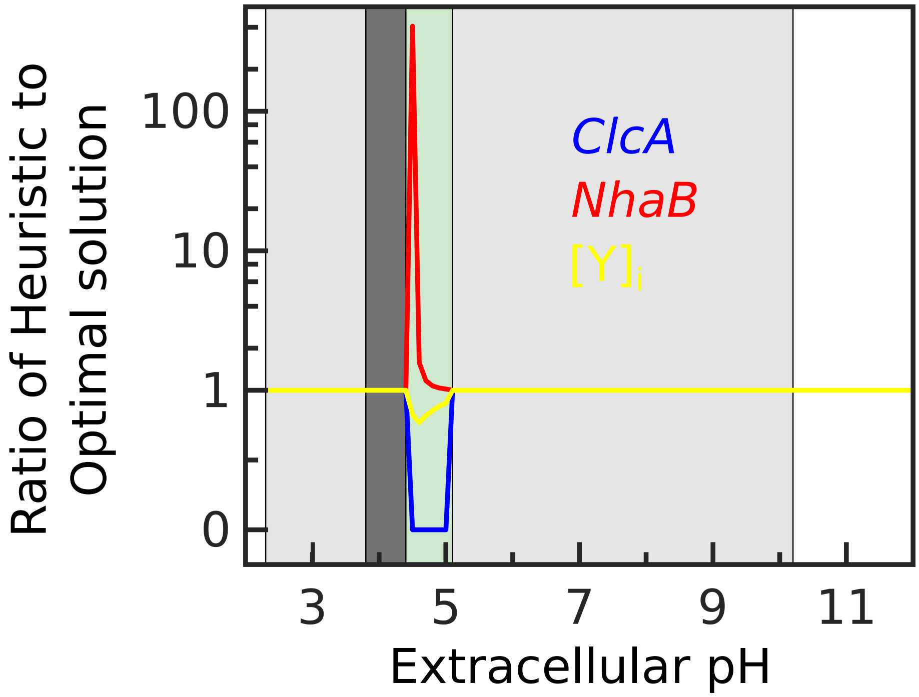


**Fig. S15. Heuristic approximation for calculating cost in Eq. S52 gives very similar results to the optimal solution, that minimizes the cost.** Both solutions set osmotic pressure to 1 atm, PMF to -170 mV and maintain pH$_i$ as close as possible to 7 (and exactly 7 in the light gray shaded area). Both solutions have identical average valency $z_Y$ and intracellular concentration $[Y]_i$, as well as forward fluxes of all antiporters apart in the shaded green area. While the heuristic solution does not allow more than one antiporter to have non zero forward fluxe, the solution minimizing the cost has in this small pH$_e$ range positive forward fluxes for both *ClcA* and *NhaB* and smaller intracellular concentration of captive charges. In this pH$_e$ range the cost of powering antiporters is low (Main text figure 4) and the cell has less incentive to minimize the cost. The average valency of captive charges and forward flux for *NhaA* are identical for all pH$_e$ for the heuristic and optimal solutions.

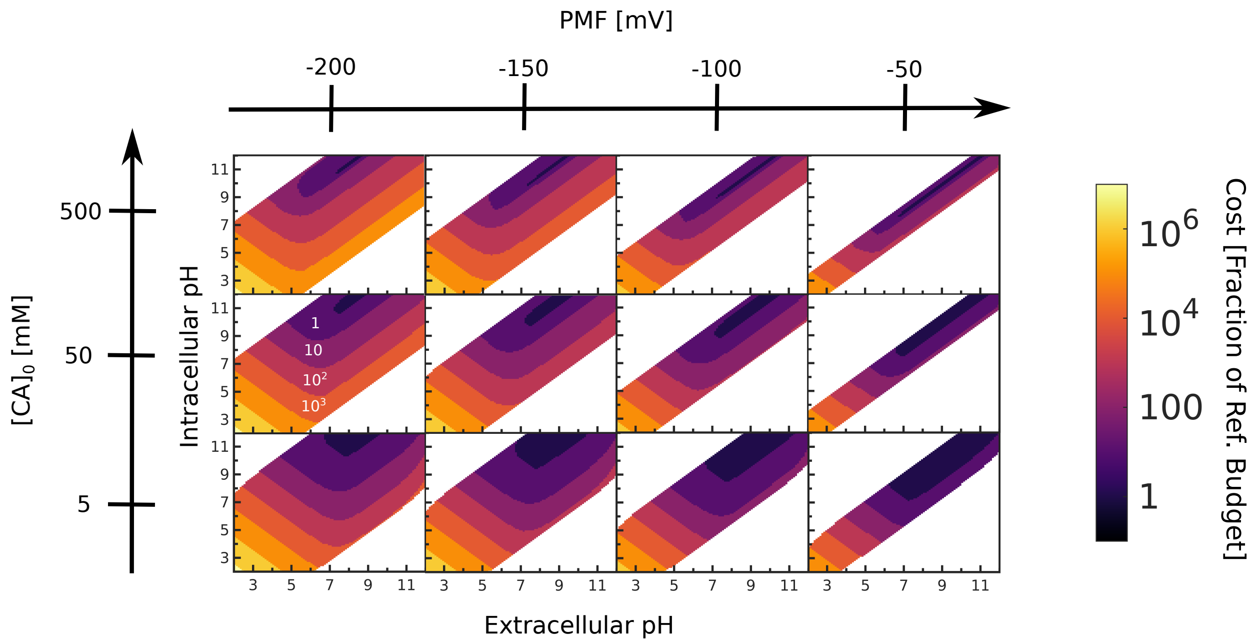


**Fig. S16. The minimal cost of maintaining particular pH$_i$ and PMF for particular extracellular environments, determined by pH$_e$ and $[\text{CA}]_0$.** We set the osmotic pressure to 1 atm and keep the average valency $-10 \leq z_Y \leq +10$, and calculate the minimal cost for a wider range of PMF values and [CA]$_0$. The cost is here estimated using the approximation for the optimal strategy calculation described above. White regions indicate PMF and pH$_i$ values that cannot be attained. White numbers give the cost value (corresponding to the colour scale).
